# Exploring the effects of genetic variation on gene regulation in cancer in the context of 3D genome structure

**DOI:** 10.1101/2020.10.06.328567

**Authors:** Noha Osman, Abd-El-Monsif Shawky, Michal Brylinski

## Abstract

Numerous genome-wide association studies (GWAS) conducted to date revealed genetic variants associated with various diseases, including breast and prostate cancers. Despite the availability of these large-scale data, relatively few variants have been functionally characterized, mainly because the majority of single-nucleotide polymorphisms (SNPs) map to the non-coding regions of the human genome. The functional characterization of these non-coding variants and the identification of their target genes remain challenging. In this communication, we explore the potential functional mechanisms of non-coding SNPs by integrating GWAS with the high-resolution chromosome conformation capture (Hi-C) data for breast and prostate cancers. We show that more genetic variants map to regulatory elements through the 3D genome structure than the 1D linear genome lacking physical chromatin interactions. Importantly, the association of enhancers, transcription factors, and their target genes with breast and prostate cancers tends to be higher when these regulatory elements are mapped to high-risk SNPs through spatial interactions compared to simply using a linear proximity. Finally, we demonstrate that topologically associating domains (TADs) carrying high-risk SNPs also contain gene regulatory elements whose association with cancer is generally higher than those belonging to control TADs containing no high-risk variants. Our results suggest that many SNPs may contribute to the cancer development by affecting the expression of certain tumor-related genes through long-range chromatin interactions with gene regulatory elements. Integrating large-scale genetic datasets with the 3D genome structure offers an attractive and unique approach to systematically investigate the functional mechanisms of genetic variants in disease risk and progression.

## Introduction

Cancer is a complex disease involving strong interactions between genetic and environmental factors, and the second leading cause of death globally [1, 2]. The dysregulation of oncogenes and/or tumor suppressor genes has an impact on cell proliferation and apoptosis in cancer pathogenesis through genetic alterations such as mutations [3, 4]. Further, the chromatin structure and regulatory elements can dysregulate gene expression subsequently leading to cancer development [5, 6]. Among different types of tumors, breast and prostate cancers create significant health problems worldwide because of their high incidence, health-related costs, and mortality rates [7, 8]. Breast cancer is the most predominant malignancy in women with a high incidence rate, prevalence, and mortality [9–11]. The extremely complex and heterogenous etiology of breast cancer, involving numerous components such as endocrine and environmental factors, other medical conditions, and genetic susceptibility [12, 13], is not yet fully understood. Prostate cancer is the second most frequent tumor in men worldwide [14]. Similar to breast cancer, it has a high genetic heritability with ethnic and geographical factors having a significant effect on the disease progression as well [15].

Genome-wide association study (GWAS) provides a systematic way to identify genetic risk factors for various diseases, including cancer [16], type 2 diabetes [17], Alzheimer’s disease [18], inflammatory bowel disease [19], and many others. The goal of GWAS is to reveal genotype-phenotype associations by detecting genomic loci that are common and low-penetrant in a specific disease state without any prior knowledge of their locations and functions [20, 21]. In the last decade, GWAS conducted on many different tumor types, including pancreatic [22], ovarian [23], lung [24], prostate [25], and breast cancer [26], identified numerous risk alleles, most of which are common and individually confer only a modest increase in disease risk. For instance, GWAS revealed 31 novel genetic susceptibility loci associated with the genetic predisposition for breast cancer [27] and 12 novel loci for prostate cancer [28]. Notably, the vast majority of genetic variants identified through GWAS (>90%) are located in the non-coding regions of the genome [29]. These variants can provide useful insights into mechanisms responsible for the development and progression of various diseases through the alteration of regulatory elements, such as transcription factors (TFs) and active enhancers, affecting the expression of certain disease-related genes [30, 31].

A number of studies investigated the downstream effects of a single-nucleotide polymorphism (SNP) in disease states [32–35]. One of the first reports was focused on a single nucleotide substitution in the promoter region of β-thalassemic globin gene expression of β-globin in patients with thalassemia [36]. Other studies investigated the effect of SNPs located in a promoter region on the promoter activity [37] as well as those located at TF binding sites affecting the binding of TFs and, subsequently, altering the gene expression [38]. Although the presence of SNPs in the non-coding regions of the genome, such as introns and intergenic regions, can alter the susceptibility to disease, the exact regulatory mechanisms of gene expression are often not fully elucidated [39, 40]. This difficulty can be attributed to the fact that SNPs may affect the expression of genes located even hundreds of kilobase pairs (kbp) away complicating the illumination of *cis*-regulatory mechanisms [31, 41]. Deciphering the effects of high-risk SNPs is not only critical to understand the molecular pathogenesis of cancer, but it can also improve cancer diagnostics and prognosis [42], and reveal potentially novel targets for pharmacotherapy [43].

Traditionally, the genome has intensively been studied as a unidimensional, linear entity often using the number of base pairs as a distance between various genomic elements. More recently, the three-dimensional (3D) structure of the genome started drawing significant attention because the regulation of gene expression and, consequently, cellular functions in physiology and disease cannot be grasped without considering the genome organization and the nuclear architecture. High-resolution chromosome conformation capture (Hi-C) is the latest variant of chromosome confirmation capture (3C) techniques developed to investigate the 3D genome structure using next-generation sequencing strategies [44]. This method enables researchers to profile pair-read contacts in all-versus-all manner in order to calculate the interaction frequency both within chromosomes (intra-chromosomal contacts) and between different chromosomes (inter-chromosomal contacts). The Hi-C resolution is determined based on the fragmentation of chromosomes and can vary from a low resolution of 1,000 kbp to as high as 5 kbp, in which each fragment comprises 5,000 base pairs [45]. Further, the genome is systematically arranged into topologically associating domains (TADs) defined as those genomic regions forming numerous self-interactions whose frequency is much higher compared to contacts with other parts of the same chromosome [46, 47]. TADs often contain multiple genes and regulatory elements, and have been shown to play a crucial role in the development of a wide array of diseases including cancer [48, 49]. Overall, the Hi-C data give invaluable insights into the 3D genome structure facilitating the identification of physical interactions among genetic variants, regulatory elements, and their corresponding target genes.

Integrating SNPs identified by GWAS with the 3D genome structure assembled from the Hi-C data offers a powerful method to investigate the genetic variation related to many human diseases [50, 51]. Indeed, several recent studies utilized Hi-C techniques in different cancer types in order to gain new insights into the effects of SNPs on regulatory elements leading to tumor progression. This approach can help identify high-risk mutations modulating gene expression in cancer by affecting regulatory elements located far away from their target genes in the linear genome [52–54]. The Hi-C data are often used in conjunction with the expression quantitative trait loci (eQTL) analysis to reveal risk loci contributing to cancer progression. For instance, the rs2981579 variant maps to the transcription start site of fibroblast growth factor receptor 2 (FGFR2) forming interaction peaks with several distal fragments [55]. These regions are located hundreds kbp from the capture region and co-localize with DNAse I hypersensitive sites, CTCF, FOXA1, GATA3, and ERα binding sites in breast cancer and normal breast epithelial cell lines. Translating these interactions helped explain the association of this SNP with FGFR2 gene regulation in breast cancer. Another example is rs4442975 associated with the susceptibility to breast cancer. According to the Hi-C data, this variant is located near a transcriptional enhancer forming physical interactions with the promoter region of insulin-like growth factor binding protein 5 (IGFBP5) [56]. IGFBP5 displays allele-specific gene expression with *g*-allele downregulating the expression of IGFBP5 leading to the increased susceptibility to breast cancer. Genetic variants are also associated with prostate cancer through long-range chromatin interactions with regulatory elements, such as the promoter regions of a specific gene (rs10486567) [51] and active enhancers regulating the expression of multiple disease-related genes (rs55958994) [57].

Although various studies were conducted to illuminate the effects of a specific genetic variation through chromatin interactions with selected gene regulatory elements, functional relationships among SNPs, regulatory elements, and disease-associated genes have not been systematically evaluated at the level of the entire human genome. In this communication, we present a large-scale analysis of the Hi-C data in the context of relationships among high-risk SNPs identified by GWAS for breast and prostate cancers, regulatory elements including TFs and enhancers, and genes associated with both diseases. The results highlight the importance of including the 3D genome structure in the investigation of the effects of genetic variation on gene regulation in cancer.

## Results

### Mapping genetic variants to regulatory elements and target genes

In this study, we compare two distinct approaches to link genetic variants highly associated with disease, with a *p*-value of ≤5×10^−8^ according to the GWAS data, to regulatory elements and their target genes (Figure 1). The first approach, schematically shown in Figure 1A, employs the unidimensional (1D) genome structure to identify those enhancers and TF binding sites located in the linear proximity up and downstream of a SNP. In this example, a TF binding site (green shape) is found downstream from a SNP (red star) and an enhancer (orange rectangle) is found upstream. For prostate cancer, the search distance is set to 5 kbp on both sides of the SNP in order to create a SNP-centered window whose size is the same as the resolution of the Hi-C data (10 kbp). Since the resolution of the Hi-C data for breast cancer used in this study is 5 kbp, we search for regulatory elements located 2.5 kbp down and 2.5 kbp upstream of a SNP. Both regulatory elements shown in Figure 1A affect the expression of their target genes, either indirectly through TF (blue teardrop) binding (genes G1-3, purple ovals) or directly (genes G4-6, yellow ovals). The search for regulatory elements in the linear proximity from 808 SNPs highly associated with breast cancer identified 12 TF binding sites affecting 8 genes for 59 SNPs and 51 enhancers affecting 33 genes for 50 SNPs. A similar search conducted for 13,447 SNPs highly associated with prostate cancer resulted in 247 TF binding sites affecting 180 genes for 986 SNPs and 3,851 enhancers affecting 613 genes for 7,641 SNPs.

**Figure 1.**
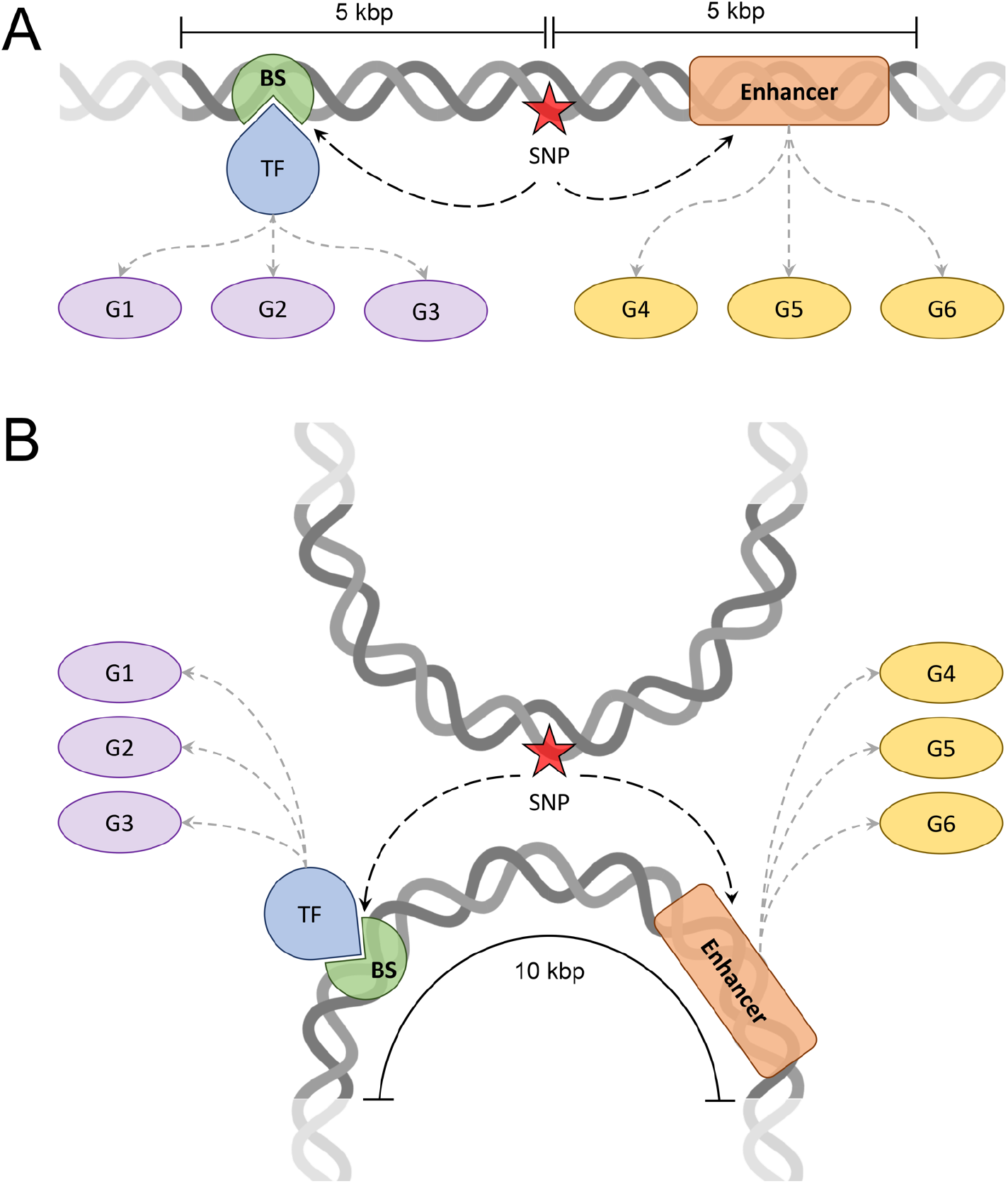
Schematic representation of the procedure to map SNPs to regulatory elements and target genes. The mapping is shown for (**A**) the 1D linear genome and (**B**) the 3D genome structure constructed at the Hi-C resolution of 10 kbp. Red stars are a SNPs highly associated with a disease at a *p*-value of ≤5×10^−8^. Regulatory elements include transcription factors (TF, blue teardrops) and their binding sites (BS, green sectors), and enhancers (orange rounded rectangles). Each regulatory element is linked to its target genes (G1-3, purple ovals for TFs and G4-6, yellow ovals for enhancers). In **A**, regulatory elements are identified within a DNA window of 10 kbp centered on the SNP, whereas in **B**, regulatory elements are identified in a DNA fragment forming a physical contact with the fragment containing the SNP.

The second approach maps SNPs highly associated with cancer at a *p*-value of ≤5×10^−8^ to regulatory elements located in the spatial proximity according to the 3D genome structure. Here, we utilize highly confident intra- and inter-chromosomal contacts obtained from the Hi-C data with a *q*-value of ≤0.05. Specifically, for each SNP, we collected those DNA fragments containing at least one regulatory element and forming physical contacts with that SNP. Next, we selected one fragment with the lowest *q*-value for a contact; in case of multiple fragments having the same lowest *q*-value for contacts, the longest-range fragment from the SNP was picked. As shown in Figure 2B, a DNA fragment containing a disease-associated SNP (red star) physically interacts with another fragment through a highly confident long-range contact. In this example, the interacting fragment contains a binding site (green shape) for a TF (blue teardrop) and an active enhancer (orange rectangle). Just as in the first approach utilizing the 1D linear genome, these elements regulate the expression of their target genes, G1-3 (purple ovals) and G4-6 (yellow ovals), respectively.

**Figure 2.**
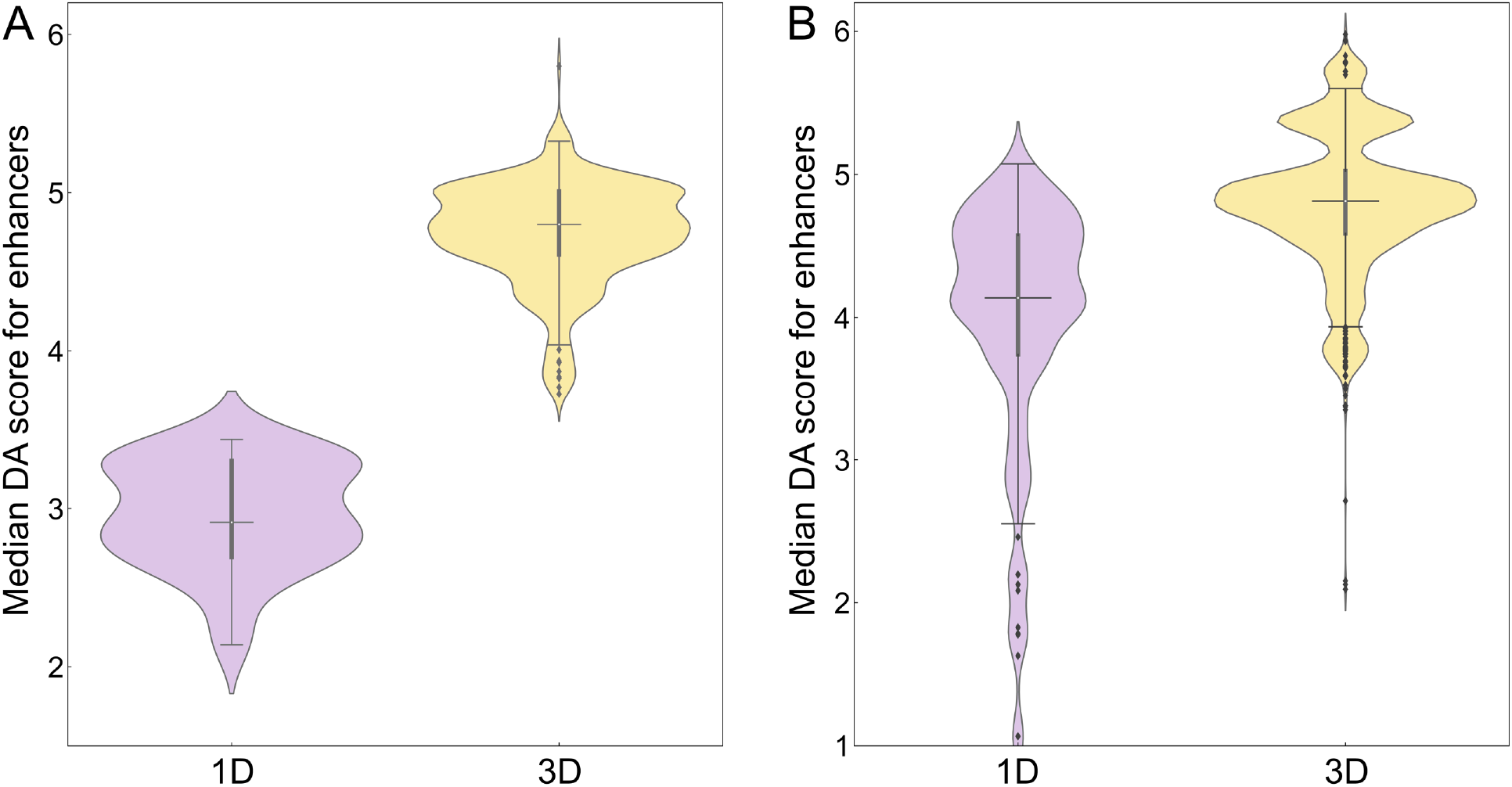
Distribution of disease-association (DA) scores for enhancers linked to SNPs. Enhancers were identified by mapping SNPs highly associated with (**A**) breast and (**B**) prostate cancer in the unidimensional (1D, purple violins) and the three-dimensional (3D, yellow violins) genome structure. In each violin, the horizontal black line is the median, the narrow gray box shows the first and third quantiles, and whiskers mark the minimum and maximum values excluding outliers, which are represented by black diamonds.

Following this procedure, we identified 19,240 chromatin fragments forming highly confident contacts with 808 SNPs associated with breast cancer. Selecting only one long-range chromatin contact per SNP with the lowest *q*-value resulted in 702 fragments containing regulatory elements. These elements include 239 enhancers having 147 target genes in contact with 702 SNPs and 83 binding sites for TFs having 70 target genes in contact with 459 SNPs. Similar to breast cancer, selecting one long-range chromatin contact per SNP with the lowest *q*-value from 1,952,907 chromatin fragments forming contacts with 13,447 SNPs associated with prostate cancer resulted in 13,429 contacts. Among these interactions, 13,410 contacts are between 13,410 SNPs and 3,585 enhancers with 747 target genes, and 1,387 contacts are between 1,387 SNPs and 324 binding sites for TFs with 190 target genes.

### Disease association of enhancers connected to genetic variants in cancer

In order to measure the relevance of those regulatory elements affected by SNPs to a disease, a series of disease association (DA) scores were computed. For each high-risk SNP, we calculated the median DA score for mapped enhancers and TFs as well as the median DA score for target genes whose expression is regulated by these elements. The number of SNPs along with quantile and inter-quantile range (IQR) values are reported in Table 1 (enhancers) and Table 2 (TFs). The distribution of DA scores for active enhancers is presented in Figure 2. Figure 2A shows that the median (2^nd^ quantile) DA score of 4.80 for enhancers linked to highly associated SNPs in breast cancer in the 3D genome structure is higher compared to 2.91 in the 1D linear genome (Table 1). Similar to breast cancer, Figure 2B and Table 1 show that the median DA score for enhancers linked to SNPs highly associated with prostate cancer is higher in 3D (4.81) than 1D (4.14). To further corroborate these results, we computed DA scores for target genes whose expression is affected by enhancers linked to genetic variants in cancer. Figure 3 and Table 1 show that the median DA scores are systematically higher in 3D compared to 1D in breast cancer (Figure 3A, 2.4 for 1D and 5.0 for 3D) and in prostate cancer (Figure 3B, 2.4 for 1D and 3.2 for 3D). In addition to higher DA scores, IQRs are typically smaller in the 3D genome structure compared to 1D (Table 1), for instance, the IQR for the enhancer DA score is 0.40 in 3D and 0.61 in 1D for breast cancer, and 0.43 in 3D and 0.83 in 1D for prostate cancer.

**Table 1.** Disease association (DA) statistics for enhancers linked to significant SNPs in breast and prostate cancer. Statistics for enhancers identified with 1D and 3D approaches include the number of SNPs used in the analysis, quantiles, and the inter-quantile range (IQR). For each cancer type, DA scores for enhancers and their target genes are reported.

| Statistic | Breast cancer |  |  |  | Prostate cancer |  |  |  |
| --- | --- | --- | --- | --- | --- | --- | --- | --- |
|  | Enhancer DA score |  | DA score for target gene |  | Enhancer DA score |  | DA score for target gene |  |
|  | 1D | 3D | 1D | 3D | 1D | 3D | 1D | 3D |
| # of SNPs | 50 | 702 | 50 | 662 | 7,641 | 13,410 | 7,213 | 12,010 |
| 1<sup>st</sup> quantile | 2.69 | 4.61 | 2.1 | 4.3 | 3.74 | 4.59 | 1.9 | 2.6 |
| 2<sup>nd</sup> quantile | 2.91 | 4.80 | 2.4 | 5.0 | 4.14 | 4.81 | 2.4 | 3.2 |
| 3<sup>rd</sup> quantile | 3.30 | 5.01 | 2.6 | 5.2 | 4.57 | 5.02 | 3.4 | 3.9 |
| IQR | 0.61 | 0.40 | 0.5 | 0.9 | 0.83 | 0.43 | 1.5 | 1.3 |

**Table 2.**
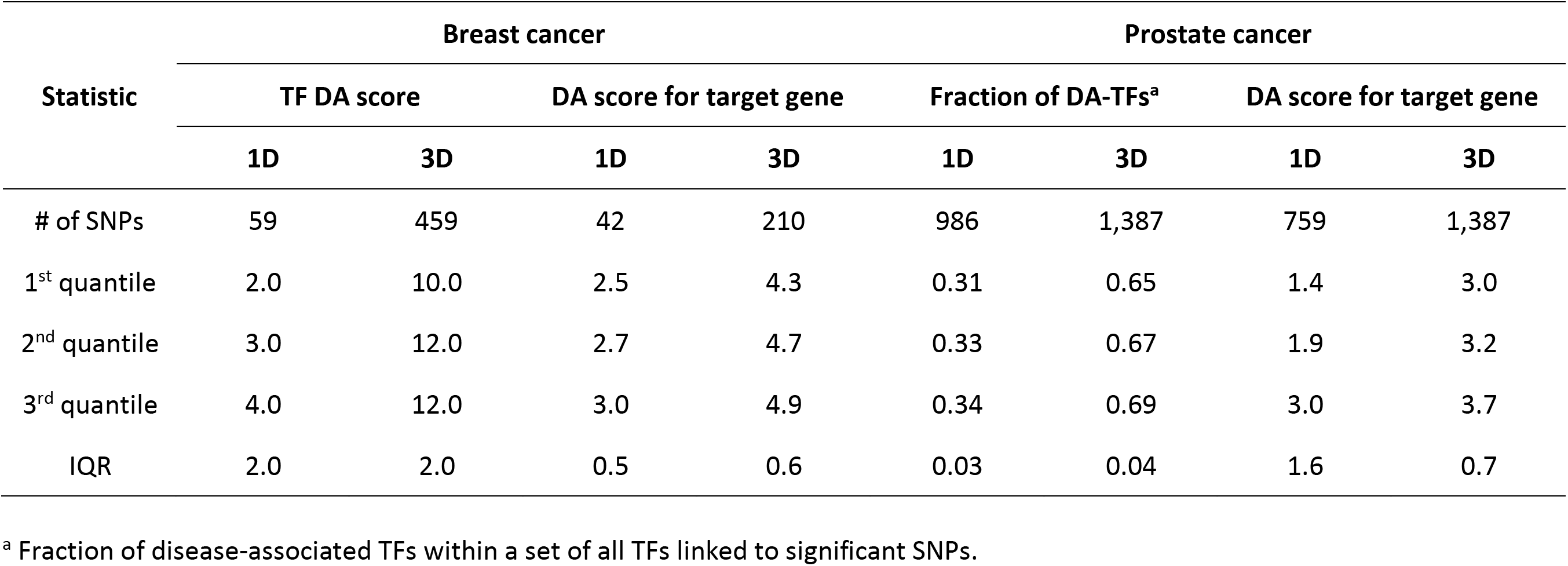
Disease association (DA) statistics for transcription factors (TFs) linked to significant SNPs in breast and prostate cancer. Statistics for enhancers identified with 1D and 3D approaches include the number of SNPs used in the analysis, quantiles, and the inter-quantile range (IQR). For each cancer type, DA scores for enhancers and their target genes are reported.

**Figure 3.**
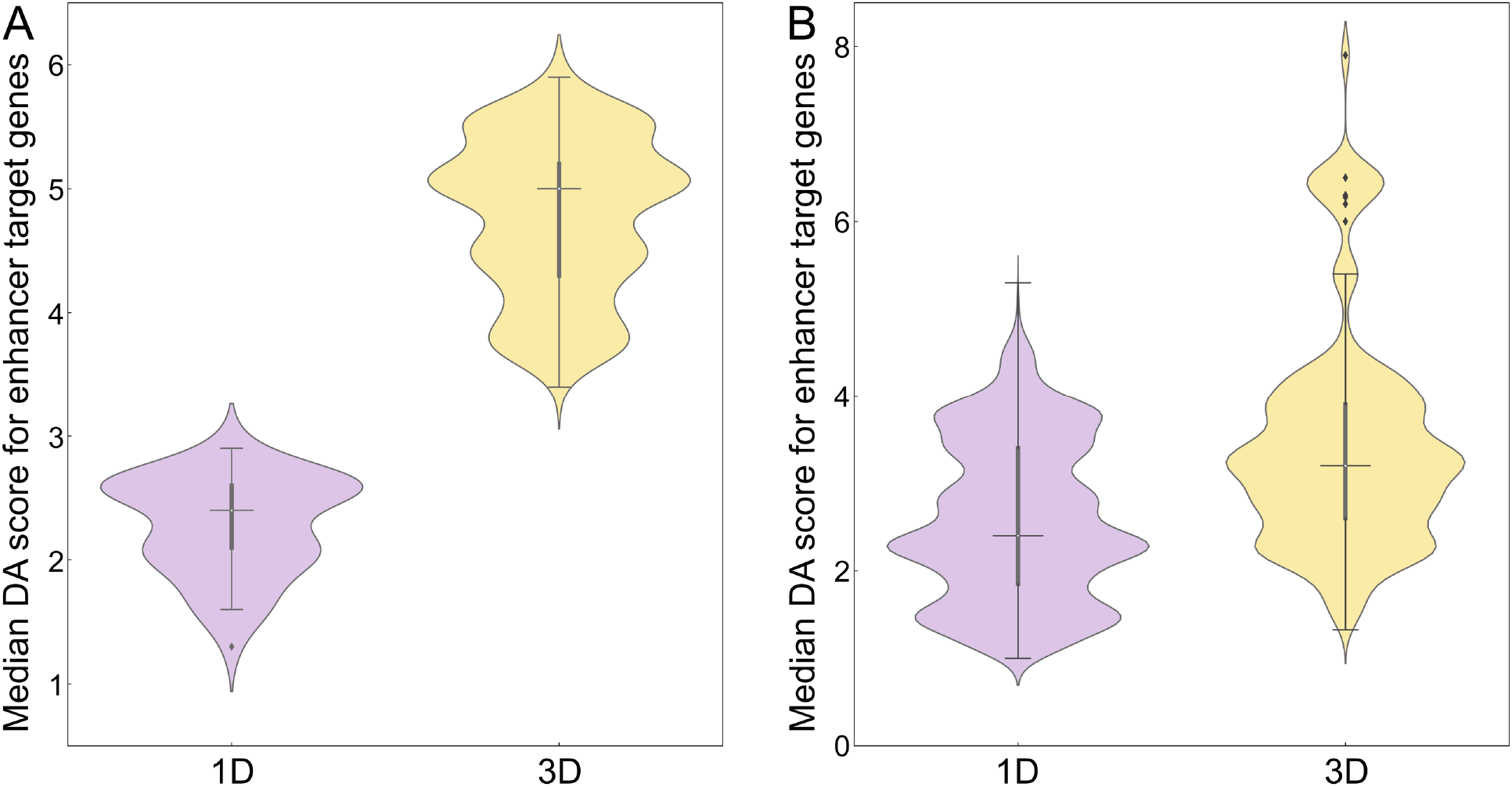
Distribution of disease-association (DA) scores for the target genes of enhancers linked to SNPs. Enhancers were first identified by mapping SNPs highly associated with (**A**) breast and (**B**) prostate cancer in the unidimensional (1D, purple violins) and the three-dimensional (3D, yellow violins) genome structure, and then linked to their target genes. In each violin, the horizontal black line is the median, the narrow gray box shows the first and third quantiles, and whiskers mark the minimum and maximum values excluding outliers, which are represented by black diamonds.

### Disease association of transcription factors connected to genetic variants in cancer

Next, we analyze the disease association of TFs linked to SNPs highly associated with cancer (Figure 4) and their target genes (Figure 5) with statistics reported in Table 2. The distribution of DA scores for TFs mapped to breast cancer is presented in Figure 4A. Here, the median DA score of 12.0 in the 3D genome structure is higher than 3.0 in 1D. In addition, Figure 5A shows that the median DA score of TF target genes is also higher in 3D (4.7) compared to 1D (2.7). In the absence of numerical DA scores for TFs linked to SNPs highly associated with prostate cancer, we conducted the analysis using the fraction of disease-associated TF (Figure 4B). On average, about two-thirds of TF mapped to SNPs in 3D are disease-associated, whereas this fraction is only one-third in 1D. Further, Figure 5B shows that the median DA score of the corresponding target genes is higher in 3D (3.2) than in 1D (1.9). In contrast to active enhancers, IQRs for TFs are similar between 1D and 3D, except for the distribution of DA scores for TF target genes in prostate cancer (1.6 and 0.7, respectively, Table 2).

**Figure 4.**
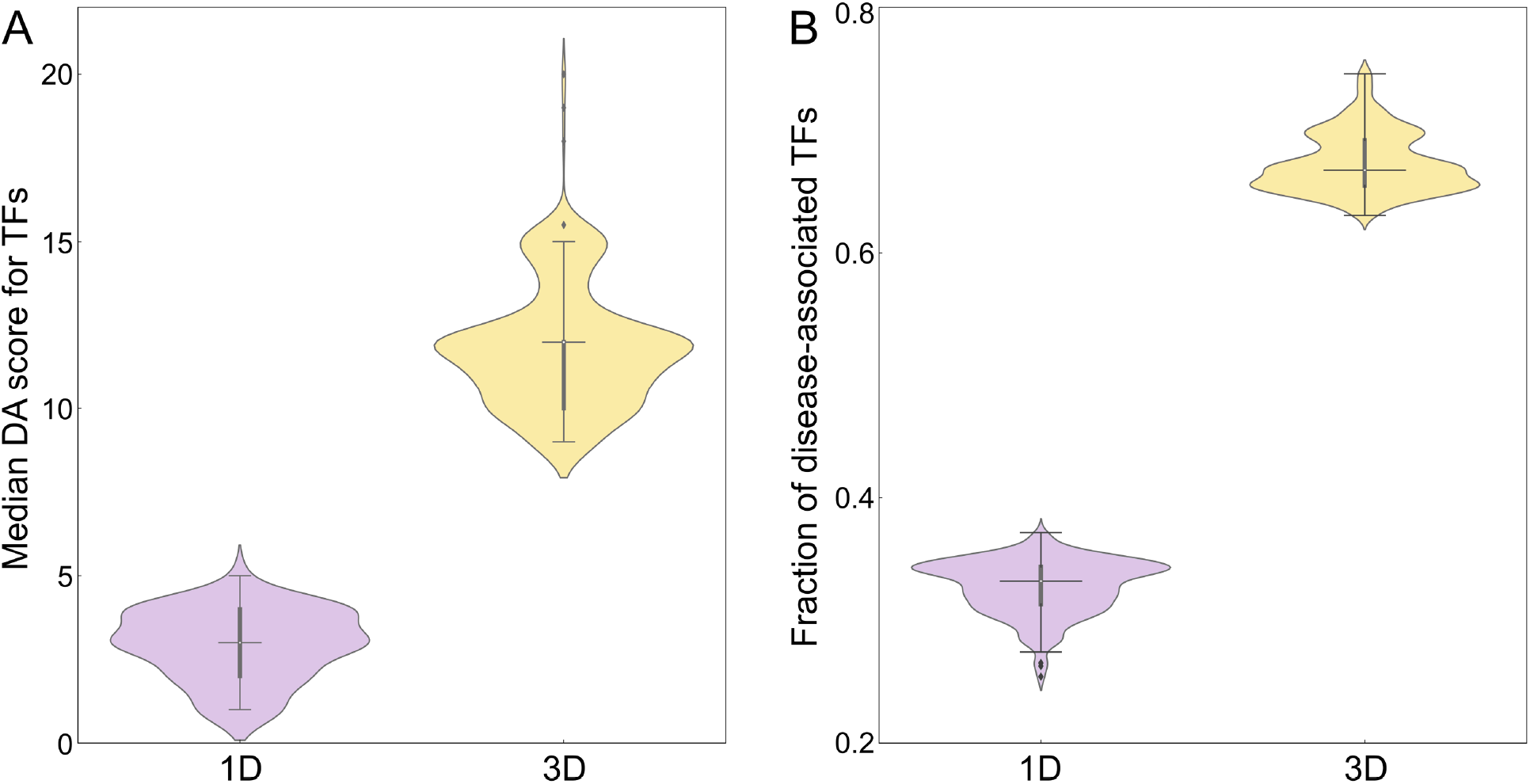
Analysis of the disease association of transcription factors (TFs) linked to SNPs. (**A**) The distribution of disease-association (DA) scores for TFs linked to SNPs highly associated with breast cancer. (**B**) The fraction of disease-associated TFs linked to SNPs highly associated with prostate cancer. TFs were identified by mapping SNPs in the unidimensional (1D, purple violins) and the three-dimensional (3D, yellow violins) genome structure. In each violin, the horizontal black line is the median, the narrow gray box shows the first and third quantiles, and whiskers mark the minimum and maximum values excluding outliers, which are represented by black diamonds.

**Figure 5.**
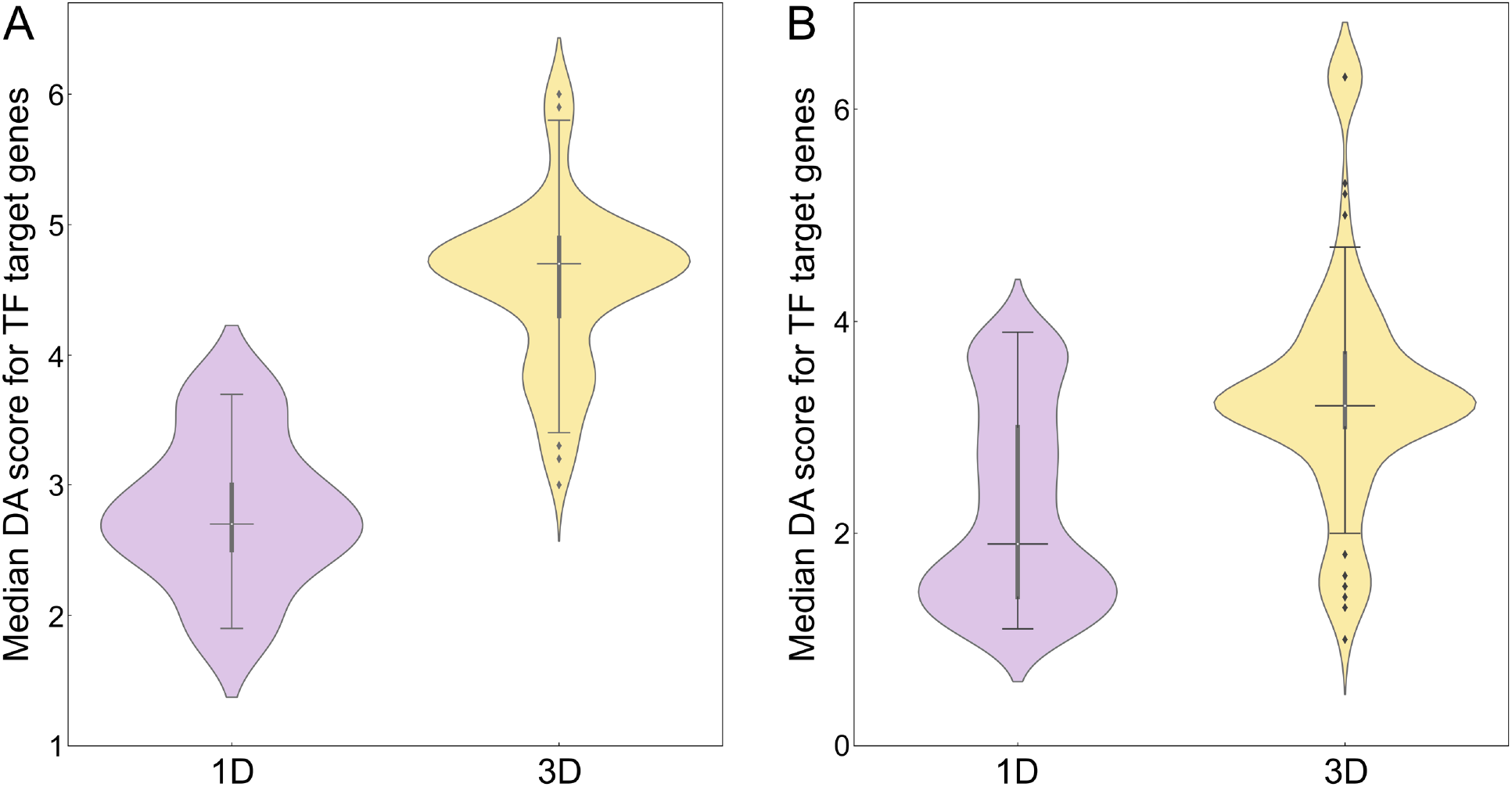
Distribution of disease-association (DA) scores for the target genes of transcription factors (TFs) linked to SNPs. TFs were first identified by mapping SNPs highly associated with (**A**) breast and (**B**) prostate cancer in the unidimensional (1D, purple violins) and the three-dimensional (3D, yellow violins) genome structure, and then mapped to their target genes. In each violin, the horizontal black line is the median, the narrow gray box shows the first and third quantiles, and whiskers mark the minimum and maximum values excluding outliers, which are represented by black diamonds.

### Examples of gene regulation through chromatin interactions in breast cancer

We present several case studies in order to illustrate the significance of the 3D genome structure in linking genetic variation to gene regulation in breast cancer (Figure 6). The locations of genomic elements discussed in this section and their association with breast cancer are provided in Supplementary Tables S1-S3. The first example is mitogen-activated protein kinase kinase kinase 1 (MAP3K1), a serine/threonine kinase known to play an important role in different functions of the cell [58, 59]. MAP3K1 can be activated by different stimuli, such as cytokines and growth factors, that constitute a complex signaling network controlling a diverse array of cellular functions [60]. In addition to numerous studies focused on somatic mutations in MAP3K1 [61–63], GWAS revealed associations between SNPs, including rs7714232 and rs16886272 regulating the expression of MAP3K1, and breast cancer [64, 65]. Further, multiple transcription factors, such as ER-α, FOXA1 and GATA3, were shown to upregulate the expression of MAP3K1 through long range chromatin interactions [64]. Figure 6A shows that rs7714232 at position Chr5:56,011,357 is in contact with a chromatin fragment containing an active enhancer 119861, and rs16886272 at position Chr5:56,067,434 is in contact with a fragment containing a putative binding site for transcription factor GATA3 predicted with a *p*-value of 2.2×10^−5^. Enhancer 119861 is associated with breast cancer at a *p*-value of 2.4×10^−5^ and GATA3 has a high disease association score of 5.9. MAP3K1, which itself has a high disease association score of 5.3, is a target gene for both regulatory elements. Thus, rs7714232 and rs16886272 may indirectly affect the expression of MAP3K1 through physical interactions with an active enhancer and a transcription factor binding site.

**Figure 6.**
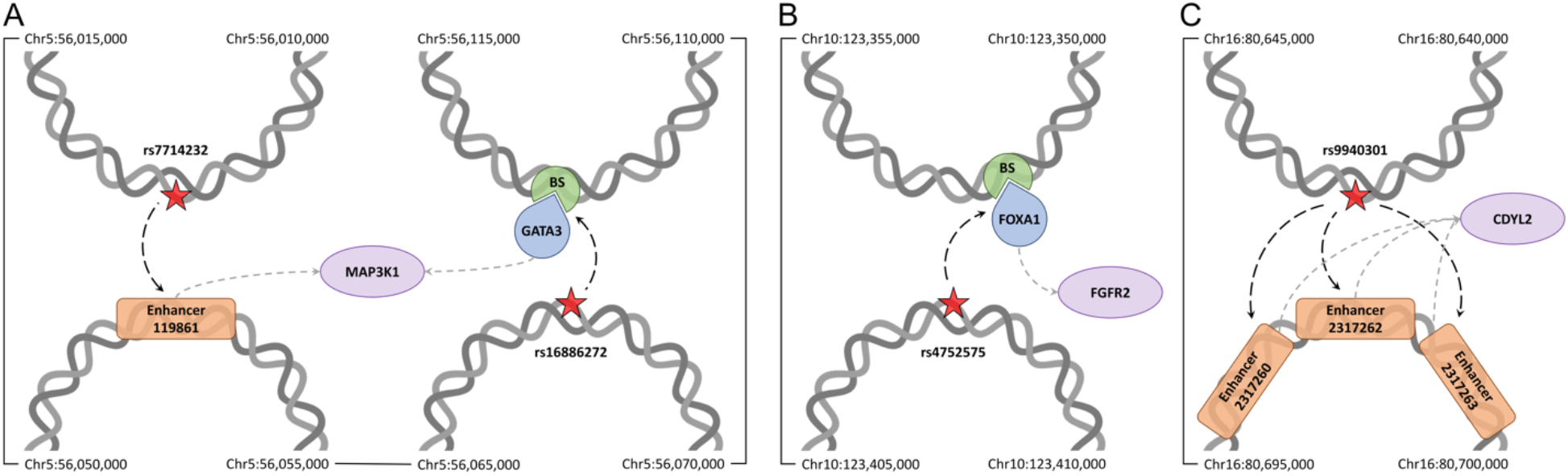
Case studies for genetic variants related to breast cancer. The possible effects of SNPs on the expression of (**A**) MAP3K1, (**B**) FGFR2, and (**C**) CDYL2 genes are presented in the context of the 3D genome structure. Red stars are SNPs highly associated with a disease at a *p*-value of ≤5×10^−8^ affecting regulatory elements through long-range physical interactions according to dashed black arrows. Dashed gray arrows link regulatory elements, including transcription factors (blue teardrops) and their binding sites (BS, green sectors), and enhancers (orange rounded rectangles) to target genes (purple ovals). Chromatin fragments from the Hi-C data (gray double helices) annotated with their start and end coordinates are connected by solid black lines showing their order in the linear genome.

Fibroblast growth factor receptor 2 (FGFR2) belonging to the receptor tyrosine kinase family mediates the cellular signaling and plays important roles in the developmental induction, cell growth and differentiation, and cell fate [66–68]. Several studies reported the association between mutations affecting FGFR2 and breast cancer [69, 70]. For example, multiple SNPs located in the second intron of FGFR2 cause the increased expression of FGFR2 linked to cancer progression [71]. Another study reported an association between the expression of FGFR2 and the number of breast tumor initiating cells [72]. GWAS data revealed the association between FGFR2 genetic variants and the risk of breast cancer [69, 73], for instance, rs4752575 was shown to alter the expression of FGFR2 leading to the increased susceptibility to breast cancer [74]. Figure 6B shows that rs4752575 located at position Chr10:123,407,187 physically interacts with a chromatin fragment containing multiple transcription factor binding sites, including a putative binding site for forkhead box protein A1 (FOXA1) predicted with a *p*-value of 4.5×10^−4^. FOXA1 is highly associated with breast cancer with a score of 5.8 and was identified as one of the master regulators of FGFR2 [75]. According to these data, we propose a new model explaining the high association of rs4752575 with breast cancer at a *p*-value of 5.5×10^−9^. Specifically, rs4752575 may dysregulate the expression of FGFR2 through the chromatin interaction with the binding site for FOXA1, a master regulator of the FGFR2 gene.

Chromodomain Y-like protein 2 (CDYL2) is a putative epigenetic factor belonging to a family of proteins characterized by the presence of N-terminal chromodomain that binds methylated histones H3K9 and H3K27 [76–78]. CDYL2 has been identified as either a tumor suppressor or oncogene depending on the cancer type [79, 80]. Moreover, CDYL2 was reported to be overexpressed in breast cancer supporting its role in disease progression [81]. The transcript variants of CDYL2 were found to be differently associated with breast cancer, suggesting a new therapeutic strategy targeting specific CDYL2 isoforms [82]. Several genetic variants related to CDYL2 have been identified by GWAS to be associated with breast cancer progression and development, including rs13329835 found in the intergenic region of CDYL2 gene [83, 84]. Another variant, rs9940301, at position Chr16:80,641,906 is strongly associated with breast cancer progression in women of African ancestry with a *p*-value of 2.0×10^−9^ [85]. According to the Hi-C data (Figure 6C), rs9940301 is in contact with a chromatin fragment containing three putative enhancers, 2317260, 2317262, and 2317263, all associated with breast cancer with a *p*-value of 9.0×10^−6^. These enhancers activate the expression of the CDYL2 gene suggesting a new association mechanism between rs9940301 and breast cancer through physical interactions with enhancers regulating CDYL2 gene expression.

### Examples of gene regulation through chromatin interactions in prostate cancer

Figure 7 presents selected cases demonstrating how the genetic variation in prostate cancer can be linked to gene regulation by analyzing the 3D genome structure. The locations of genomic elements discussed in this section and their association with prostate cancer are provided in Supplementary Tables S1-S3. Androgen receptor (AR) is a master regulator belonging to the nuclear receptor superfamily [86]. It acts as a transcription factor binding a specific ligand molecule to control the expression of targeted genes [87]. Prostate function primarily depends on the androgen signaling axis through the regulation of AR target genes [88]. AR is highly associated with prostate cancer with a score of 8.0, which is consistent with observations that it is often overexpressed in prostate cancer [89] and mutations in the AR gene are present in a large population of castration-resistant prostate cancer patients [90, 91]. The GWAS data revealed that numerous SNPs near the AR locus are associated with prostate cancer [92, 93]. For instance, rs6152 is located in the first exon of the AR gene and it is associated with a susceptibility to prostate cancer at a *p*-value of 1.5×10^−12^ [94, 95]. We also found that rs6152 at position ChrX:66,765,627 forms a contact with a chromatin fragment containing an active enhancer 2765787 associated with prostate cancer at a *p*-value of 1.6×10^−2^ and affecting the expression of AR (Figure 7A). Thus, the physical interaction between rs6152 and an enhancer may play a role in the regulation of AR gene expression during prostate cancer progression.

**Figure 7.**
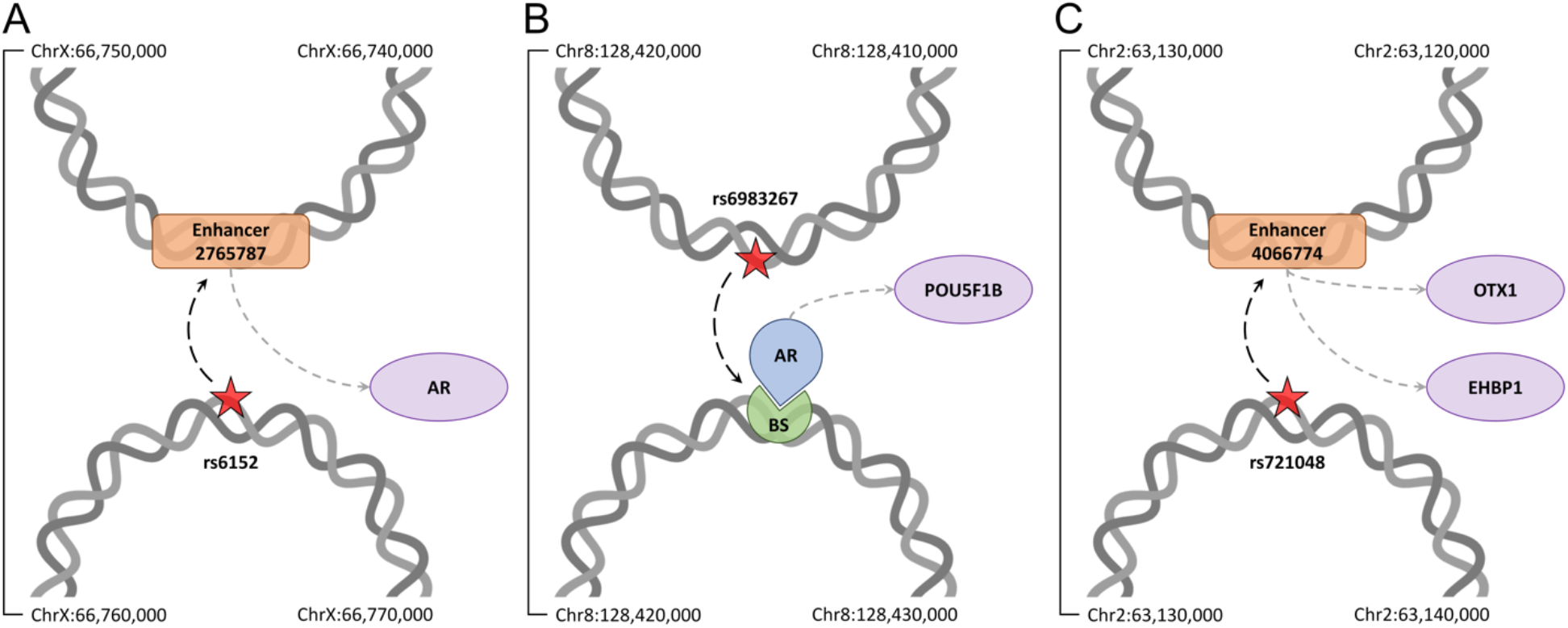
Case studies for genetic variants related to prostate cancer. The possible effects of SNPs on the expression of (**A**) AR, (**B**) POU5F1B, and (**C**) OXT1 and EHBP1 genes are presented in the context of the 3D genome structure. Red stars are SNPs highly associated with a disease at a *p*-value of ≤5×10^−8^ affecting regulatory elements through long-range physical interactions according to dashed black arrows. Dashed gray arrows link regulatory elements, including transcription factors (blue teardrops) and their binding sites (BS, green sectors), and enhancers (orange rounded rectangles) to target genes (purple ovals). Chromatin fragments from the Hi-C data (gray double helices) annotated with their start and end coordinates are connected by solid black lines showing their order in the linear genome.

Octamer-binding transcription factor 4 (OCT4), a member of the POU domain-containing family of transcription factors, is expressed in embryonic and adult stem cells [96]. Although six different pseudogenes identified for OCT4 are not expressed, these elements are believed to play a role in the regulation of OCT4 expression [97, 98]. Interestingly, two of these OCT4 pseudogenes, POU5F1P5 and POU5F1B, were found to be transcribed in cancer cells [99]. POU5F1B was shown to be overexpressed in gastric cancer and its knockdown confirmed a role for POU5F1B in the promotion of tumor cell growth [100]. Further, the methylation level near the POU5F1B gene [101] and the genetic variation around that region [102] were found to be associated with the prostate cancer risk. For instance, rs6983267 was reported to be in linkage disequilibrium with the open-reading frame of the POU5F1B gene among people of European ancestry and associated with the expression of POU5F1B in prostate of white subjects [103]. Located at position Chr8:128,413,305, rs6983267 is associated with prostate cancer at a *p*-value of 2.8×10^−141^. Figure 7B shows that it is also in contact with a chromatin fragment containing multiple putative transcription factor binding sites including a confidently predicted binding site for AR with a *p*-value of 3.5×10^−4^, which regulates the expression of POU5F1B. According to these findings, rs6983267 may play a role in regulating PO5F1B expression by affecting the binding of AR to its binding site.

EH domain-binding protein 1 (EHBP1) gene encodes Eps15 homology domain binding protein playing a role in endocytic trafficking [104]. Recently, GWAS reported the association of a genetic variant rs721048, located within one of the introns of the EHBP1 gene, and the susceptibility to prostate cancer [28, 105] with a *p*-value of 5.0×10^−22^. Interestingly, Figure 7C shows that rs721048 at position Chr2:63,131,731 forms a contact with a chromatin fragment containing an active enhancer 406774 associated with prostate cancer at a *p*-value of 1.5×10^−4^ that regulates the expression of EHBP1. Further, enhancer 406774 also regulates the expression of orthodenticle homeobox 1 (OTX1), a transcription factor playing a critical role in multiple developmental processes, such as the neuronal differentiation [106]. Several studies reported the hypermethylation of the OTX1 gene promoter region in non-small lung cancer [107, 108] and an altered OTX1 expression in medulloblastoma and other brain tumors [109]. It is important to note that the expression of OTX1 is also associated with prostate cancer risk [110]. Further, OTX1 is one of several transcription factors involved in tumor-specific enhancer networks and it was found to be linked to active enhancers in prostate adenocarcinoma [111]. Our data suggest that rs721048 may be associated with prostate cancer through the disruption of the mechanism of action of certain tumor-specific enhancers causing the dysregulation of the expression of OTX1 and EHBP1 genes.

### Mapping genetic variants to topologically associating domains

Next, TADs were identified from the Hi-C data and all regulatory elements and SNPs were mapped to these domains as shown in Figure 8. Based on the presence of SNPs highly associated with cancer, the resulting TADs are divided into two groups, TADs containing no SNPs (control, Figure 8A) and TADs containing at least one SNP with a p-value of ≤5×10^−8^ according to the GWAS data (SNP-rich, Figure 8B). Specifically, we identified the total of 21,157 TADs from the Hi-C data for breast cancer, including 30 TADs enriched with disease-associated SNPs. Among 30 SNP-rich TADs, 26 also contain enhancers (477 in total) and 17 contain TF binding sites (36 in total). The control dataset for breast cancer comprises 16,473 TADs containing 259,780 enhancers and 10,463 TADs containing 23,890 binding sites for TFs. In addition, the total of 17,435 TADs were detected from the Hi-C data for prostate cancer, including 304 TADs enriched with disease-associated SNPs; 291 of these SNP-rich TADs contain enhancers (7,940 in total) and 241 contain TF binding sites (686 in total). The control dataset for prostate cancer comprises 13,587 TADs containing 250,789 enhancers and 10,070 TADs containing 22,562 binding sites for TFs.

**Figure 8.**
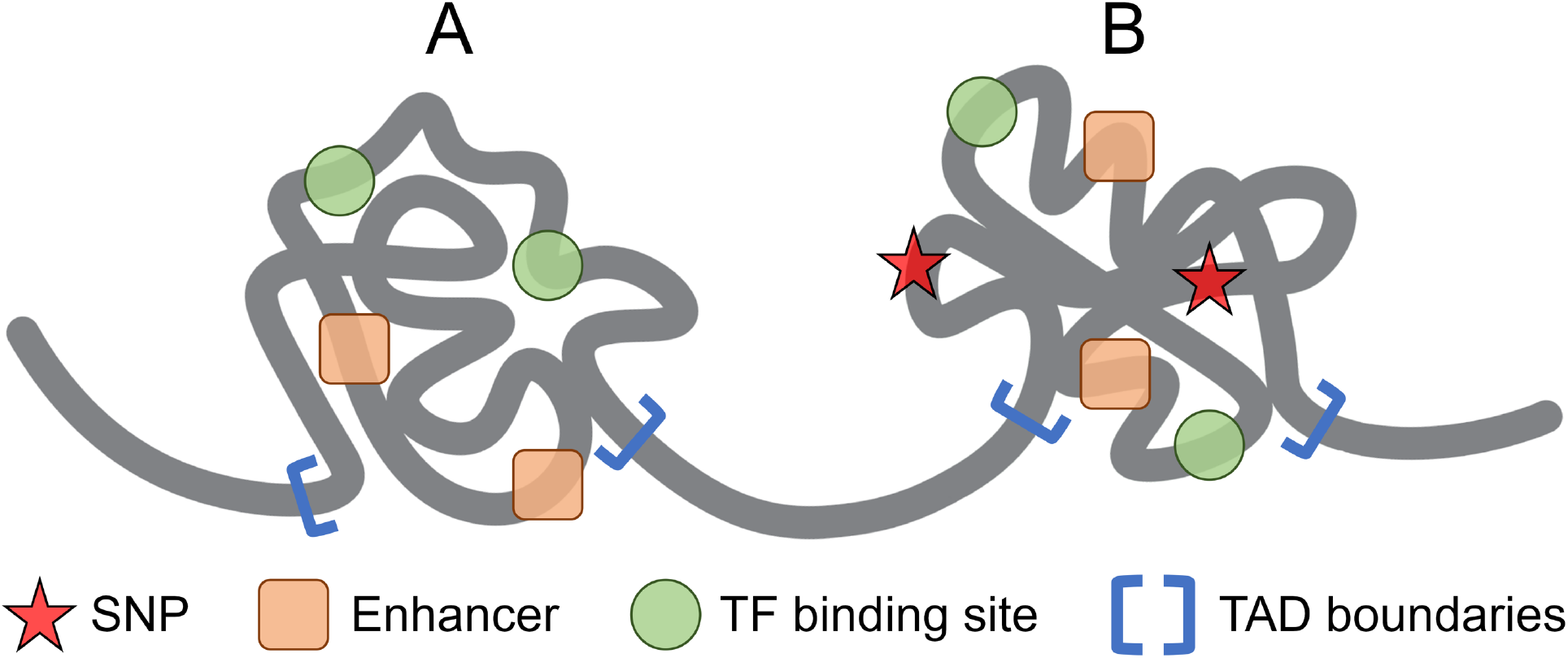
Schematic representation of topologically associating domains (TADs). Red stars are SNPs highly associated with a disease at a *p*-value of ≤5×10^−8^, orange rounded squares represent enhancers, green circles represent transcription factors, and blue square brackets show the location of TAD boundaries. TADs are divided into two groups, (**A**) control TADs containing no highly associated SNPs and (**B**) SNP-rich TADs carrying at least one highly associated variant.

We first take a glance at selected genetic variants highly associated with breast (4 SNPs) and prostate (3 SNPs) cancers discussed above in order to determine whether gene regulatory elements located in their spatial proximity according to the Hi-C data reside in the same TAD. Interestingly, this holds true in almost all cases. Both variants rs7714232 and rs16886272 associated with breast cancer, enhancer 119861, and a binding site for transcription factor GATA3 are located in TAD 6450 whose boundaries are Chr5:56,010,000 – Chr5:56,140,000. Further, variant rs9940301 along with all three enhancers, 2317260, 2317262, and 2317263, reside in the same TAD 17447 (Chr16:80,630,000 – Chr16:80,950,000). Variant rs4752575 and FOXA1 binding site belong to neighboring TADs 12498 (Chr10:123,360,000 – Chr10:123,460,000) and 12497 (Chr10:123,330,000 – Chr10:123,360,000), respectively. In prostate cancer, TAD 1830 (Chr2:63,010,000 – Chr2:63,160,000) contains both rs721048 and enhancer 406774, TAD 8618 (Chr8:128,410,000 – Chr8:128,490,000) contains both rs6983267 and AR binding site, and TAD 16953 (ChrX:66,530,000 – ChrX:67,270,000) contains both rs6152 and enhancer 2765787. On that account, we expect TADs enriched in disease-associated SNPs to also contain those gene regulatory elements having higher disease association compared to control TADs, which is investigated in the following section.

### Association of SNPs and regulatory elements with cancer in the context of TADs

In order to verify the assertion that those TADs carrying genetic variants highly associated with cancer also contain gene regulatory elements whose disease association is high, we first calculated DA scores for enhancers located within control and SNP-rich TADs. The distribution of DA scores is shown in Figure 9 with the corresponding statistics reported in Table 3. Compared to a median DA score of 4.75 for control TADs in breast cancer, SNP-rich TADs contain enhancers with a higher median DA score of 5.66 (Figure 9A). A median DA score of 5.66 for enhancers located in SNP-rich TADs is higher than a value of 4.69 for those belonging to control TADs in prostate cancer as well (Figure 9B). Similar to enhancers, Figure 10 shows the distribution of DA scores computed for TFs residing in SNP-rich and control TADs with the corresponding statistics reported in Table 3. In breast cancer, the median DA score of 11.5 for TFs located in control TADs is lower than a value of 17.0 for those present in SNP-rich TADs, whereas in prostate cancer, the fraction of disease-associated TFs in SNP-rich TADs is twice as high as in control TADs (Figure 10B). Further, Table 3 shows that IQRs for enhancers and TFs are very similar between SNP-rich and control TADs in both cancers. These results demonstrate that disease-associated genetic variants and gene regulatory elements indeed tend co-localize within certain TADs identified based on the 3D genome structure.

**Figure 9.**
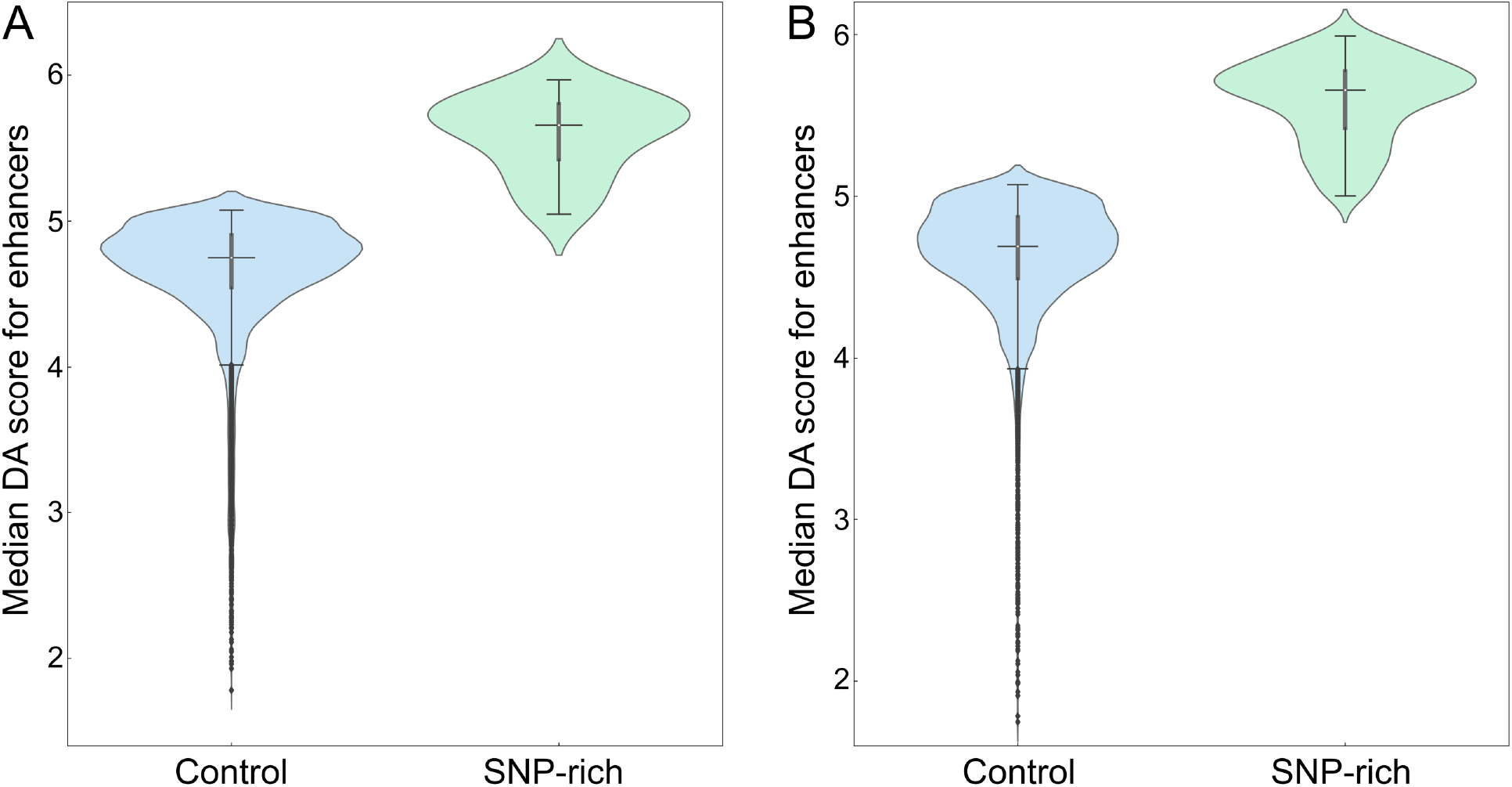
Distribution of disease-association (DA) scores for enhancers within TADs. DA scores against (**A**) breast and (**B**) prostate cancer are calculated for enhancers present in control (blue violins) and SNP-rich (green violins) TADs. In each violin, the horizontal black line is the median, the narrow gray box shows the first and third quantiles, and whiskers mark the minimum and maximum values excluding outliers, which are represented by black diamonds.

**Table 3.** Disease association (DA) statistics for enhancers and transcription factors (TFs) within TADs in breast and prostate cancer. TADs are divided into two groups, containing no SNPs with a significant association to disease (control) and those containing at least one SNP with a significant disease association (SNP-rich). Statistics include the number of TADs used in the analysis, quantiles, and the inter-quantile range (IQR).

| Statistic | Breast cancer |  |  |  | Prostate cancer |  |  |  |
| --- | --- | --- | --- | --- | --- | --- | --- | --- |
|  | Enhancer DA score |  | TF DA score |  | Enhancer DA score |  | Fraction of DA-TFs<sup>a</sup> |  |
|  | Control | SNP-rich | Control | SNP-rich | Control | SNP-rich | Control | SNP-rich |
| # of TADs | 16,473 | 26 | 10,463 | 17 | 13,587 | 291 | 10,070 | 241 |
| 1<sup>st</sup> quantile | 4.55 | 5.42 | 10.5 | 17.0 | 4.50 | 5.42 | 0.32 | 0.66 |
| 2<sup>nd</sup> quantile | 4.75 | 5.66 | 11.5 | 17.0 | 4.69 | 5.66 | 0.33 | 0.67 |
| 3<sup>rd</sup> quantile | 4.90 | 5.80 | 12.5 | 18.0 | 4.87 | 5.77 | 0.35 | 0.69 |
| IQR | 0.35 | 0.38 | 2.0 | 1.0 | 0.37 | 0.35 | 0.03 | 0.03 |

<sup>a</sup> Fraction of disease-associated TFs within TADs.
Fraction of disease-associated TFs

**Figure 10.**
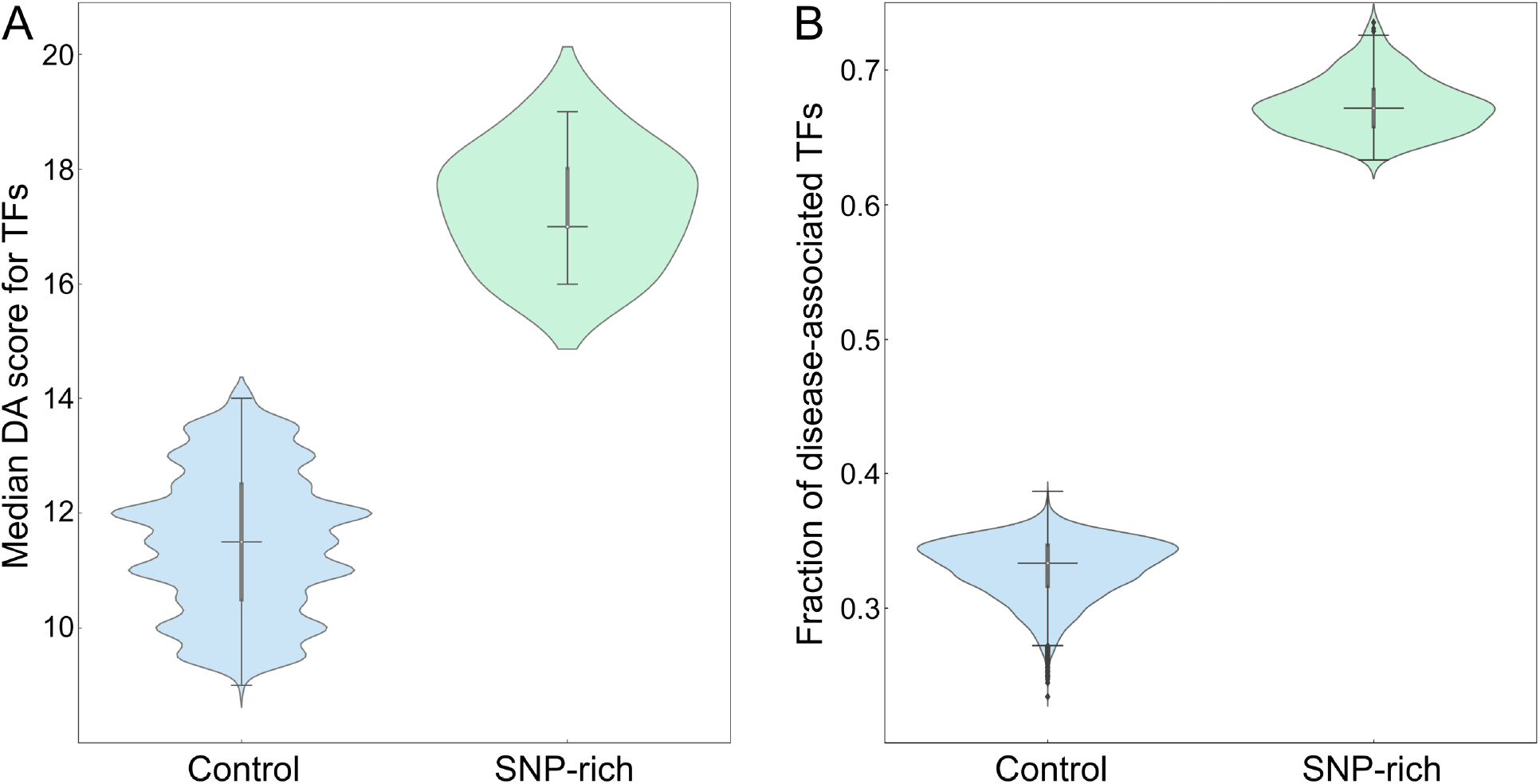
Analysis of the disease association of transcription factors (TFs) within TADs. (**A**) The distribution of disease-association (DA) scores against breast cancer and (**B**) the fraction of TFs associated with prostate cancer. Blue violins show the distribution of DA scores and the fraction within control TADs, whereas green violins show the distribution within SNP-rich TADs. In each violin, the horizontal black line is the median, the narrow gray box shows the first and third quantiles, and whiskers mark the minimum and maximum values excluding outliers, which are represented by black diamonds.

## Discussion

Although a large number of genetic variants associated with cancer have been identified by GWAS, the exact mechanisms by which these mutations affect the phenotype have not yet been fully elucidated. A significant challenge in explaining the functional mechanisms of SNPs in cancer initiation and progression arises from the fact that the vast majority of disease-associated variants are located within the non-coding regions of the genome. Non-coding SNPs are thought to exert their pathological effects by altering gene regulation rather than directly affecting the sequence, structure, and function of gene products. Accumulated data on the 3D genome structure collected from Hi-C experiments offer a unique opportunity to investigate the effects of genetic variation, particularly those located in the non-coding complement of the human genome, on gene regulation leading to cancer pathophenotypes. In this communication, we integrated the large-scale data provided by GWAS with the information on chromatin structure and interactions in breast and prostate cancers in order to systematically evaluate the effects of SNPs on gene regulatory mechanisms. We are specifically interested in the comparison of this 3D method utilizing the Hi-C data to a 1D proximity approach assuming that genetic variants affect regulatory elements located down- and up-stream in the linear DNA.

Focusing on SNPs highly associated with breast and prostate cancers, we conducted a comprehensive analysis of their relationships to various active enhancers and binding sites for transcription factors regulating the expression of their target genes. Considering a vast body of evidence supporting the association of many regulatory elements with cancer phenotypes, we propose that genetic variants forming physical contacts with these elements may contribute to the development and progression of disease by altering the expression levels of cancer-related genes. There are several benefits of including the Hi-C data in the analysis of the effects of genetic variation on gene expression. The 3D genome structure allows for a more efficient mapping between variants and regulatory elements compared to the unidimensional genome lacking physical chromatin interactions [112, 113]. Importantly, the disease association scores of enhancers and transcription factors mapped to cancer-associated SNPs through chromatin contacts are consistently higher compared to those identified using a linear DNA model. This also holds true for the target genes of these regulatory elements. Our results are in line with numerous studies demonstrating that the 3D structure of the genome, which facilitates certain DNA-DNA interactions regulating gene expression [114], can effectively be used to reveal the functional mechanisms of genetic variation as well as candidate genes related to cancer [115].

Finally, we investigated the relationship among genetic variants, regulatory elements, and their target genes in the context of TADs, relatively small, compact, and self-interacting genomic regions [116]. Interestingly, we found that those TADs carrying cancer-associated SNPs also contain enhancers and binding sites for transcription factors whose disease association is generally higher compared to regulatory elements located in control TADs devoid of high-risk SNPs. Overall, these data can help reveal hot spots within the human genome linked to disease. Integrating the chromatin structure with the genomic data is a promising approach to study how the spatial organization of the genome affects gene regulation through physical interactions between various genomic regions. This technique also holds promise to investigate subtle differences between the genetic makeup of individuals leading to varying levels of disease risk and progression. Our work provides a new perspective for investigating the effects of genetic variation on gene regulation in cancer through the large-scale analysis of long-range chromatin interactions shaping the 3D genome structure.

## Methods

### Hi-C data

Hi-C data at 5 kbp resolution collected for the human mammary epithelial cell line were downloaded from the Gene Expression Omnibus database (GEO accession: GSE63525) [117]. The Hi-C data at 10 kbp resolution collected for the human normal prostate cell line were downloaded from the Gene Expression Omnibus database (GEO accession: GSM3564252) [118]. FitHiC2 was applied to intra- and inter-chromosomal contacts to exclude low-confidence interactions [119] keeping only those contacts whose *q*-values (*p*-values corrected with the false discovery rate) is ≤0.05 [120].

### Genome-wide association studies

In this study, we used the GWAS data for breast cancer generated for 122,977 cases and 105,974 controls of European ancestry and 14,068 cases and 13,104 of East Asian ancestry [121], and for prostate cancer generated for 46,939 cases and 27,910 controls of European ancestry [122]. We first identified SNPs having OncoArray accession numbers resulting in 568,712 SNPs for breast cancer and 13,502,794 SNPs for prostate cancer, and then we selected 808 (breast cancer) and 13,447 (prostate cancer) highly associated SNPs with a *p*-value of ≤5×10^−8^ [123].

### Gene regulatory elements

Data on active enhancers, including their genomic location, target genes, and disease association scores were obtained from the Human Enhancer Disease Database (HEDD) database [124]. HEDD provides comprehensive information for about 2.8 million human enhancers identified by ENCODE, FANTOM5 and RoadMap with disease association scores based on enhancer-gene and gene-disease associations. In this study, we used 262,490 enhancers related to breast cancer and 267,453 enhancers related to prostate cancer. The data on transcription factors were obtained from the TF2DNA database containing 1,306 TFs, 19,190 target genes, and 24,634,759 binding sites [125]. From these data, we selected 164 TFs associated with breast cancer [126] and 612 TFs associated with prostate cancer [127]. Disease association scores for enhancer and TF target genes were collected from the DISEASES database providing evidence on disease-gene associations computed from automatic text mining, manually curated literature, cancer mutation data, and genome-wide association studies [128]. We identified the total of 12,846 genes having an association score to breast cancer and 10,032 genes having an association score to prostate cancer.

### Genome-wide mapping of genetic variants to regulatory elements

In the unidimensional approach, SNPs highly associated with breast and prostate cancer were mapped to enhancers and TF binding sites in the linear proximity using a 5 kbp window (2.5 kbp up and 2.5 kbp downstream from the SNP) for breast cancer and a 10 kbp window (5 kbp up and 5 kbp downstream from the SNP) for prostate cancer. These window sizes were selected to match the resolution of the Hi-C data. For each SNP, we then calculated the median association score of mapped enhancers and TFs as well as their target genes with the exception of TFs in prostate cancer, for which we computed the fraction of disease-associated TFs. In the three-dimensional approach, each highly associated SNP was mapped to regulatory elements present in a chromatin fragment forming the most confident contacts with the lowest *q*-value. When multiple fragments have the same lowest *q*-value, we selected the longest-range interaction. This way, the same number of chromatin fragments are utilized by both 1D and 3D approaches. The median association scores of regulatory elements and their target genes mapped through physical interactions were calculated in a similar manner as in the 1D analysis.

### Topologically associating domains

TADs were identified from intra-chromosomal contacts for each chromosome with the TopDom software [129]. For each TAD, we calculated the median association scores for regulatory elements present in that domain as well as for their target genes. TADs were then divided into two groups, SNP-rich carrying at least one genetic variant highly associated with cancer at a *p*-value of ≤5×10^−8^ and control TADs containing no highly associated SNPs.

## Data availability

The generated data is available through the Open Science Framework (OSF) project at https://osf.io/4xwc6/.

## Acknowledgements

This work has been supported in part by the National Institute of General Medical Sciences of the National Institutes of Health under Award Number R35GM119524. Portions of this research were conducted with high-performance computational resources provided by Louisiana State University.

## Author contributions

NO and AS collected and processed the data. NO conducted data analyses. NO and AS selected case studies. NO drafted the manuscript. MB coordinated the project and prepared the final version of the manuscript.

## Additional information

Supplementary information is available for this paper.

## References

1. Knox, S.S., From ‘omics’ to complex disease: a systems biology approach to gene-environment interactions in cancer. Cancer Cell Int, 2010. 10: p. 11.

2. Nagai, H. and Y.H. Kim, Cancer prevention from the perspective of global cancer burden patterns. J Thorac Dis, 2017. 9(3): p. 448–451.

3. Vogelstein, B. and K.W. Kinzler, Cancer genes and the pathways they control. Nat Med, 2004. 10(8): p. 789–99.

4. Zhu, K., et al., Oncogenes and tumor suppressor genes: comparative genomics and network perspectives. BMC Genomics, 2015. 16 Suppl 7: p. S8.

5. See, Y.X., B.Z. Wang, and M.J. Fullwood, Chromatin interactions and regulatory elements in cancer: From bench to bedside. Trends Genet, 2019. 35(2): p. 145–158.

6. Flavahan, W.A., E. Gaskell, and B.E. Bernstein, Epigenetic plasticity and the hallmarks of cancer. Science, 2017. 357(6348).

7. Rawla, P., Epidemiology of prostate cancer. World J Oncol, 2019. 10(2): p. 63–89.

8. Sharma, R., Breast cancer incidence, mortality and mortality-to-incidence ratio (MIR) are associated with human development, 1990-2016: evidence from Global Burden of Disease Study 2016. Breast Cancer, 2019. 26(4): p. 428–445.

9. Eccles, S.A., et al., Critical research gaps and translational priorities for the successful prevention and treatment of breast cancer. Breast Cancer Res, 2013. 15(5): p. R92.

10. Lima, Z.S., et al., Recent advances of therapeutic targets based on the molecular signature in breast cancer: genetic mutations and implications for current treatment paradigms. J Hematol Oncol, 2019. 12(1): p. 38.

11. Narod, S.A., V. Giannakeas, and V. Sopik, Time to death in breast cancer patients as an indicator of treatment response. Breast Cancer Res Treat, 2018. 172(3): p. 659–669.

12. Anothaisintawee, T., et al., Risk factors of breast cancer: a systematic review and meta-analysis. Asia Pac J Public Health, 2013. 25(5): p. 368–87.

13. Pharoah, P.D., et al., Polygenic susceptibility to breast cancer and implications for prevention. Nat Genet, 2002. 31(1): p. 33–6.

14. Bell, K.J., et al., Prevalence of incidental prostate cancer: A systematic review of autopsy studies. Int J Cancer, 2015. 137(7): p. 1749–57.

15. Mucci, L.A., et al., Familial risk and heritability of cancer among twins in Nordic countries. JAMA, 2016. 315(1): p. 68–76.

16. Sud, A., B. Kinnersley, and R.S. Houlston, Genome-wide association studies of cancer: current insights and future perspectives. Nat Rev Cancer, 2017. 17(11): p. 692–704.

17. Xue, A., et al., Genome-wide association analyses identify 143 risk variants and putative regulatory mechanisms for type 2 diabetes. Nat Commun, 2018. 9(1): p. 2941.

18. Nazarian, A., A.I. Yashin, and A.M. Kulminski, Genome-wide analysis of genetic predisposition to Alzheimer’s disease and related sex disparities. Alzheimers Res Ther, 2019. 11(1): p. 5.

19. Cho, J.H., The genetics and immunopathogenesis of inflammatory bowel disease. Nat Rev Immunol, 2008. 8(6): p. 458–66.

20. Kim, M.S., et al., Genetic disease risks can be misestimated across global populations. Genome Biol, 2018. 19(1): p. 179.

21. Tam, V., et al., Benefits and limitations of genome-wide association studies. Nat Rev Genet, 2019. 20(8): p. 467–484.

22. Klein, A.P., et al., Genome-wide meta-analysis identifies five new susceptibility loci for pancreatic cancer. Nat Commun, 2018. 9(1): p. 556.

23. Yodsurang, V., et al., Genome-wide association study (GWAS) of ovarian cancer in Japanese predicted regulatory variants in 22q13.1. PLoS One, 2018. 13(12): p. e0209096.

24. McKay, J.D., et al., Large-scale association analysis identifies new lung cancer susceptibility loci and heterogeneity in genetic susceptibility across histological subtypes. Nat Genet, 2017. 49(7): p. 1126–1132.

25. Benafif, S., et al., A review of prostate cancer genome-wide association studies (GWAS). Cancer Epidemiol Biomarkers Prev, 2018. 27(8): p. 845–857.

26. Ferreira, M.A., et al., Genome-wide association and transcriptome studies identify target genes and risk loci for breast cancer. Nat Commun, 2019. 10(1): p. 1741.

27. Shu, X., et al., Identification of novel breast cancer susceptibility loci in meta-analyses conducted among Asian and European descendants. Nat Commun, 2020. 11(1): p. 1217.

28. Takata, R., et al., 12 new susceptibility loci for prostate cancer identified by genome-wide association study in Japanese population. Nat Commun, 2019. 10(1): p. 4422.

29. Abecasis, G.R., et al., A map of human genome variation from population-scale sequencing. Nature, 2010. 467(7319): p. 1061–73.

30. Hrdlickova, B., et al., Genetic variation in the non-coding genome: Involvement of micro-RNAs and long non-coding RNAs in disease. Biochim Biophys Acta, 2014. 1842(10): p. 1910–1922.

31. Madelaine, R., et al., A screen for deeply conserved non-coding GWAS SNPs uncovers a MIR-9-2 functional mutation associated to retinal vasculature defects in human. Nucleic Acids Res, 2018. 46(7): p. 3517–3531.

32. Huo, Y., et al., Functional genomics reveal gene regulatory mechanisms underlying schizophrenia risk. Nat Commun, 2019. 10(1): p. 670.

33. Rojano, E., et al., Regulatory variants: from detection to predicting impact. Brief Bioinform, 2019. 20(5): p. 1639–1654.

34. Wilk, G. and R. Braun, regQTLs: Single nucleotide polymorphisms that modulate microRNA regulation of gene expression in tumors. PLoS Genet, 2018. 14(12): p. e1007837.

35. Li, M.J., et al., Exploring the function of genetic variants in the non-coding genomic regions: approaches for identifying human regulatory variants affecting gene expression. Brief Bioinform, 2015. 16(3): p. 393–412.

36. Orkin, S.H., S.E. Antonarakis, and H.H. Kazazian, Base substitution at position −88 in a beta-thalassemic globin gene. Further Evidence for the Role of Distal Promoter Element ACACCC. J. Biol. Chem., 1984. 259: p. 8679–8681.

37. Horn, S., et al., TERT promoter mutations in familial and sporadic melanoma. Science, 2013. 339(6122): p. 959–61.

38. Bond, G.L., et al., A single nucleotide polymorphism in the MDM2 promoter attenuates the p53 tumor suppressor pathway and accelerates tumor formation in humans. Cell, 2004. 119(5): p. 591–602.

39. Khurana, E., et al., Role of non-coding sequence variants in cancer. Nat Rev Genet, 2016. 17(2): p. 93–108.

40. van Arensbergen, J., et al., High-throughput identification of human SNPs affecting regulatory element activity. Nat Genet, 2019. 51(7): p. 1160–1169.

41. Fagny, M., et al., Nongenic cancer-risk SNPs affect oncogenes, tumour-suppressor genes, and immune function. Br J Cancer, 2020. 122(4): p. 569–577.

42. Deng, N., et al., Single nucleotide polymorphisms and cancer susceptibility. Oncotarget, 2017. 8(66): p. 110635–110649.

43. Palmirotta, R., et al., SNPs in predicting clinical efficacy and toxicity of chemotherapy: walking through the quicksand. Oncotarget, 2018. 9(38): p. 25355–25382.

44. Lieberman-Aiden, E., et al., Comprehensive mapping of long-range interactions reveals folding principles of the human genome. Science, 2009. 326(5950): p. 289–93.

45. Belton, J.M., et al., Hi-C: a comprehensive technique to capture the conformation of genomes. Methods, 2012. 58(3): p. 268–76.

46. Dixon, J.R., D.U. Gorkin, and B. Ren, Chromatin domains: The unit of chromosome organization. Mol Cell, 2016. 62(5): p. 668–80.

47. Pombo, A. and N. Dillon, Three-dimensional genome architecture: players and mechanisms. Nat Rev Mol Cell Biol, 2015. 16(4): p. 245–57.

48. Galupa, R. and E. Heard, Topologically associating domains in chromosome architecture and gene regulatory landscapes during development, disease, and evolution. Cold Spring Harb Symp Quant Biol, 2017. 82: p. 267–278.

49. Valton, A.L. and J. Dekker, TAD disruption as oncogenic driver. Curr Opin Genet Dev, 2016. 36: p. 34–40.

50. Beesley, J., et al., Chromatin interactome mapping at 139 independent breast cancer risk signals. Genome Biol, 2020. 21(1): p. 8.

51. Luo, Z., et al., A prostate cancer risk element functions as a repressive loop that regulates HOXA13. Cell Rep, 2017. 21(6): p. 1411–1417.

52. Jager, R., et al., Capture Hi-C identifies the chromatin interactome of colorectal cancer risk loci. Nat Commun, 2015. 6: p. 6178.

53. Ji, P., et al., Systematic analyses of genetic variants in chromatin interaction regions identified four novel lung cancer susceptibility loci. J Cancer, 2020. 11(5): p. 1075–1081.

54. O’Mara, T.A., et al., Analysis of promoter-associated chromatin interactions reveals biologically relevant candidate target genes at endometrial cancer risk loci. Cancers (Basel), 2019. 11(10).

55. Baxter, J.S., et al., Capture Hi-C identifies putative target genes at 33 breast cancer risk loci. Nat Commun, 2018. 9(1): p. 1028.

56. Ghoussaini, M., et al., Evidence that breast cancer risk at the 2q35 locus is mediated through IGFBP5 regulation. Nat Commun, 2014. 4: p. 4999.

57. Qian, Y., et al., The prostate cancer risk variant rs55958994 regulates multiple gene expression through extreme long-range chromatin interaction to control tumor progression. Sci Adv, 2019. 5(7): p. eaaw6710.

58. Kyriakis, J.M. and J. Avruch, Mammalian mitogen-activated protein kinase signal transduction pathways activated by stress and inflammation. Physiol Rev, 2001. 81(2): p. 807–69.

59. Uhlik, M.T., et al., Wiring diagrams of MAPK regulation by MEKK1, 2, and 3. Biochem Cell Biol, 2004. 82(6): p. 658–63.

60. Cuevas, B.D., A.N. Abell, and G.L. Johnson, Role of mitogen-activated protein kinase kinase kinases in signal integration. Oncogene, 2007. 26(22): p. 3159–71.

61. Avivar-Valderas, A., et al., Functional significance of co-occurring mutations in PIK3CA and MAP3K1 in breast cancer. Oncotarget, 2018. 9(30): p. 21444–21458.

62. Michaut, M., et al., Integration of genomic, transcriptomic and proteomic data identifies two biologically distinct subtypes of invasive lobular breast cancer. Sci Rep, 2016. 6: p. 18517.

63. Pereira, B., et al., The somatic mutation profiles of 2,433 breast cancers refines their genomic and transcriptomic landscapes. Nat Commun, 2016. 7: p. 11479.

64. Glubb, D.M., et al., Fine-scale mapping of the 5q11.2 breast cancer locus reveals at least three independent risk variants regulating MAP3K1. Am J Hum Genet, 2015. 96(1): p. 5–20.

65. Mocellin, S., et al., Breast cancer susceptibility: an integrative analysis of genomic data. bioRxiv, 2018: p. 279984.

66. Eswarakumar, V.P., I. Lax, and J. Schlessinger, Cellular signaling by fibroblast growth factor receptors. Cytokine Growth Factor Rev, 2005. 16(2): p. 139–49.

67. Touat, M., et al., Targeting FGFR signaling in cancer. Clin Cancer Res, 2015. 21(12): p. 2684–94.

68. Turner, N. and R. Grose, Fibroblast growth factor signalling: from development to cancer. Nat Rev Cancer, 2010. 10(2): p. 116–29.

69. Hunter, D.J., et al., A genome-wide association study identifies alleles in FGFR2 associated with risk of sporadic postmenopausal breast cancer. Nat Genet, 2007. 39(7): p. 870–4.

70. Udler, M.S., et al., FGFR2 variants and breast cancer risk: fine-scale mapping using African American studies and analysis of chromatin conformation. Hum Mol Genet, 2009. 18(9): p. 1692–703.

71. Meyer, K.B., et al., Allele-specific up-regulation of FGFR2 increases susceptibility to breast cancer. PLoS Biol, 2008. 6(5): p. e108.

72. Kim, S., et al., FGFR2 promotes breast tumorigenicity through maintenance of breast tumor-initiating cells. PLoS One, 2013. 8(1): p. e51671.

73. Barnholtz-Sloan, J.S., et al., FGFR2 and other loci identified in genome-wide association studies are associated with breast cancer in African-American and younger women. Carcinogenesis, 2010. 31(8): p. 1417–23.

74. Meyer, K.B., et al., Fine-scale mapping of the FGFR2 breast cancer risk locus: putative functional variants differentially bind FOXA1 and E2F1. Am J Hum Genet, 2013. 93(6): p. 1046–60.

75. Fletcher, M.N., et al., Master regulators of FGFR2 signalling and breast cancer risk. Nat Commun, 2013. 4: p. 2464.

76. Dorus, S., et al., The CDY-related gene family: coordinated evolution in copy number, expression profile and protein sequence. Hum Mol Genet, 2003. 12(14): p. 1643–50.

77. Fischle, W., et al., Specificity of the chromodomain Y chromosome family of chromodomains for lysine-methylated ARK(S/T) motifs. J Biol Chem, 2008. 283(28): p. 19626–35.

78. Franz, H., et al., Multimerization and H3K9me3 binding are required for CDYL1b heterochromatin association. J Biol Chem, 2009. 284(50): p. 35049–59.

79. Mulligan, P., et al., CDYL bridges REST and histone methyltransferases for gene repression and suppression of cellular transformation. Mol Cell, 2008. 32(5): p. 718–26.

80. Wu, H., et al., Short-Form CDYLb but not long-form CDYLa functions cooperatively with histone methyltransferase G9a in hepatocellular carcinomas. Genes Chromosomes Cancer, 2013. 52(7): p. 644–55.

81. Siouda, M., et al., CDYL2 epigenetically regulates MIR124 to control NF-kappaB/STAT3-dependent breast cancer cell plasticity. iScience, 2020. 23(6): p. 101141.

82. Yang, L.F., et al., Discrete functional and mechanistic roles of chromodomain Y-like 2 (CDYL2) transcript variants in breast cancer growth and metastasis. Theranostics, 2020. 10(12): p. 5242–5258.

83. Ghoussaini, M., P.D.P. Pharoah, and D.F. Easton, Inherited genetic susceptibility to breast cancer: the beginning of the end or the end of the beginning? Am J Pathol, 2013. 183(4): p. 1038–1051.

84. Michailidou, K., et al., Large-scale genotyping identifies 41 new loci associated with breast cancer risk. Nat Genet, 2013. 45(4): p. 353–61, 361e1–2.

85. Feng, Y., et al., Characterizing genetic susceptibility to breast cancer in women of African ancestry. Cancer Epidemiol Biomarkers Prev, 2017. 26(7): p. 1016–1026.

86. Mangelsdorf, D.J., et al., The nuclear receptor superfamily: the second decade. Cell, 1995. 83(6): p. 835–9.

87. Chang, C., et al., Androgen receptor: an overview. Crit Rev Eukaryot Gene Expr, 1995. 5(2): p. 97–125.

88. Buchanan, G., et al., Contribution of the androgen receptor to prostate cancer predisposition and progression. Cancer Metastasis Rev, 2001. 20(3-4): p. 207–23.

89. LaTulippe, E., et al., Comprehensive gene expression analysis of prostate cancer reveals distinct transcriptional programs associated with metastatic disease. Cancer Res, 2002. 62(15): p. 4499–506.

90. Grasso, C.S., et al., The mutational landscape of lethal castration-resistant prostate cancer. Nature, 2012. 487(7406): p. 239–43.

91. Waltering, K.K., A. Urbanucci, and T. Visakorpi, Androgen receptor (AR) aberrations in castration-resistant prostate cancer. Mol Cell Endocrinol, 2012. 360(1-2): p. 38–43.

92. Eeles, R.A., et al., Identification of 23 new prostate cancer susceptibility loci using the iCOGS custom genotyping array. Nat Genet, 2013. 45(4): p. 385–91, 391e1–2.

93. Kote-Jarai, Z., et al., Seven prostate cancer susceptibility loci identified by a multi-stage genome-wide association study. Nat Genet, 2011. 43(8): p. 785–91.

94. Cucchiara, V., et al., Association between Rs6152 polymorphism in the androgen receptor gene and disease aggressiveness in a prospective cohort of prostate cancer patients undergoing radical prostatectomy. Journal of Urology, 2018. 199(4): p. E372–E372.

95. Lu, J. and M. Danielsen, A Stu I polymorphism in the human androgen receptor gene (AR). Clin Genet, 1996. 49(6): p. 323–4.

96. Pochampally, R.R., et al., Serum deprivation of human marrow stromal cells (hMSCs) selects for a subpopulation of early progenitor cells with enhanced expression of OCT-4 and other embryonic genes. Blood, 2004. 103(5): p. 1647–52.

97. Bai, M., et al., OCT4 pseudogene 5 upregulates OCT4 expression to promote proliferation by competing with miR-145 in endometrial carcinoma. Oncol Rep, 2015. 33(4): p. 1745–52.

98. Pain, D., et al., Multiple retropseudogenes from pluripotent cell-specific gene expression indicates a potential signature for novel gene identification. J Biol Chem, 2005. 280(8): p. 6265–8.

99. Suo, G., et al., Oct4 pseudogenes are transcribed in cancers. Biochem Biophys Res Commun, 2005. 337(4): p. 1047–51.

100. Hayashi, H., et al., The OCT4 pseudogene POU5F1B is amplified and promotes an aggressive phenotype in gastric cancer. Oncogene, 2015. 34(2): p. 199–208.

101. Barry, K.H., et al., Prospective study of DNA methylation at chromosome 8q24 in peripheral blood and prostate cancer risk. Br J Cancer, 2017. 116(11): p. 1470–1479.

102. Yeager, M., et al., Genome-wide association study of prostate cancer identifies a second risk locus at 8q24. Nat Genet, 2007. 39(5): p. 645–9.

103. Breyer, J.P., et al., An expressed retrogene of the master embryonic stem cell gene POU5F1 is associated with prostate cancer susceptibility. Am J Hum Genet, 2014. 94(3): p. 395–404.

104. Guilherme, A., et al., EHD2 and the novel EH domain binding protein EHBP1 couple endocytosis to the actin cytoskeleton. J Biol Chem, 2004. 279(11): p. 10593–605.

105. Ao, X., et al., Association between EHBP1 rs721048(A>G) polymorphism and prostate cancer susceptibility: a meta-analysis of 17 studies involving 150,678 subjects. Onco Targets Ther, 2015. 8: p. 1671–80.

106. Huang, B., et al., OTX1 regulates cell cycle progression of neural progenitors in the developing cerebral cortex. J Biol Chem, 2018. 293(6): p. 2137–2148.

107. Jin, M., et al., Different histological types of non-small cell lung cancer have distinct folate and DNA methylation levels. Cancer Sci, 2009. 100(12): p. 2325–30.

108. Kalari, S. and G.P. Pfeifer, Identification of driver and passenger DNA methylation in cancer by epigenomic analysis. Adv Genet, 2010. 70: p. 277–308.

109. Omodei, D., et al., Expression of the brain transcription factor OTX1 occurs in a subset of normal germinal-center B cells and in aggressive Non-Hodgkin Lymphoma. Am J Pathol, 2009. 175(6): p. 2609–17.

110. Hazelett, D.J., et al., Comprehensive functional annotation of 77 prostate cancer risk loci. PLoS Genet, 2014. 10(1): p. e1004102.

111. Rhie, S.K., et al., Identification of activated enhancers and linked transcription factors in breast, prostate, and kidney tumors by tracing enhancer networks using epigenetic traits. Epigenetics Chromatin, 2016. 9: p. 50.

112. Mishra, A. and R.D. Hawkins, Three-dimensional genome architecture and emerging technologies: looping in disease. Genome Med, 2017. 9(1): p. 87.

113. Schoenfelder, S., et al., Promoter capture Hi-C: High-resolution, genome-wide profiling of promoter interactions. J Vis Exp, 2018(136).

114. Ron, G., et al., Promoter-enhancer interactions identified from Hi-C data using probabilistic models and hierarchical topological domains. Nat Commun, 2017. 8(1): p. 2237.

115. Du, M., et al., Chromatin interactions and candidate genes at ten prostate cancer risk loci. Sci Rep, 2016. 6: p. 23202.

116. Rowley, M.J. and V.G. Corces, The three-dimensional genome: principles and roles of long-distance interactions. Curr Opin Cell Biol, 2016. 40: p. 8–14.

117. Rao, S.S., et al., A 3D map of the human genome at kilobase resolution reveals principles of chromatin looping. Cell, 2014. 159(7): p. 1665–80.

118. Rhie, S.K., et al., A high-resolution 3D epigenomic map reveals insights into the creation of the prostate cancer transcriptome. Nat Commun, 2019. 10(1): p. 4154.

119. Kaul, A., S. Bhattacharyya, and F. Ay, Identifying statistically significant chromatin contacts from Hi-C data with FitHiC2. Nat Protoc, 2020. 15(3): p. 991–1012.

120. Chakraborty, A. and F. Ay, Identification of copy number variations and translocations in cancer cells from Hi-C data. Bioinformatics, 2018. 34(2): p. 338–345.

121. Michailidou, K., et al., Association analysis identifies 65 new breast cancer risk loci. Nature, 2017. 551(7678): p. 92–94.

122. Schumacher, F.R., et al., Association analyses of more than 140,000 men identify 63 new prostate cancer susceptibility loci. Nat Genet, 2018. 50(7): p. 928–936.

123. Pe’er, I., et al., Estimation of the multiple testing burden for genomewide association studies of nearly all common variants. Genet Epidemiol, 2008. 32(4): p. 381–5.

124. Wang, Z., et al., HEDD: Human Enhancer Disease Database. Nucleic Acids Res, 2018. 46(D1): p. D113–D120.

125. Pujato, M., et al., Prediction of DNA binding motifs from 3D models of transcription factors; identifying TLX3 regulated genes. Nucleic Acids Res, 2014. 42(22): p. 13500–12.

126. Ji, Z., et al., Genome-scale identification of transcription factors that mediate an inflammatory network during breast cellular transformation. Nat Commun, 2018. 9(1): p. 2068.

127. Dhingra, P., et al., Identification of novel prostate cancer drivers using RegNetDriver: a framework for integration of genetic and epigenetic alterations with tissue-specific regulatory network. Genome Biol, 2017. 18(1): p. 141.

128. Pletscher-Frankild, S., et al., DISEASES: text mining and data integration of disease-gene associations. Methods, 2015. 74: p. 83–9.

129. Shin, H., et al., TopDom: an efficient and deterministic method for identifying topological domains in genomes. Nucleic Acids Res, 2016. 44(7): p. e70.

