## Supplemental Tables S1-S3 for "Exploring the effects of genetic variation on gene regulation in cancer in the context of 3D genome structure" (from another version of this preprint: version 2, 2020-11-05)

**Table S1.** Genetic variants selected to illustrate the significance of the 3D genome structure in gene regulation in breast cancer.

| Genetic variant | Genomic position | TAD[a] | Disease association[b] |
|---|---|---|---|
| *Breast cancer* | | | |
| rs7714232 | Chr5:56,011,357 | 6450 | $9.1\times10^{-13}$ |
| rs16886272 | Chr5:56,067,434 | | $4.7\times10^{-9}$ |
| rs4752575 | Chr10:123,407,187 | 12498 | $5.5\times10^{-9}$ |
| rs9940301 | Chr16:80,641,906 | 17447 | $2.0\times10^{-9}$ |
| *Prostate cancer* | | | |
| rs6152 | ChrX:66,765,627 | 16953 | $1.5\times10^{-12}$ |
| rs6983267 | Chr8:128,413,305 | 8618 | $2.8\times10^{-141}$ |
| rs721048 | Chr2:63,131,731 | 1830 | $5.0\times10^{-22}$ |

[a] ID of topologically associating domain, [b] *p*-value from genome-wide association studies.

**Table S2.** Enhancers forming physical contacts with genetic variants listed in Table S1 according to the 3D genome structure.

| Enhancer ID | Genomic position | TAD[a] | Disease association[b] | Target genes | DA-score[c] |
|---|---|---|---|---|---|
| | | *Breast cancer* | | | |
| 119861 | Chr5:56,052,902-56,053,241 | 6450 | $2.5\times10^{-5}$ | MAP3K1 | 5.3 |
| 2317260 | Chr16:80,698,090-80,698,168 | | | | |
| 2317262 | Chr16:80,698,782-80,698,981 | 17447 | $9.0\times10^{-6}$ | CDYL2 | 2.0 |
| 2317263 | Chr16:80,699,105-80,699,234 | | | | |
| | | *Prostate cancer* | | | |
| 2765787 | ChrX:66,744,392-66,744,590 | 16953 | $1.6\times10^{-2}$ | AR | 8.0 |
| 406774 | Chr2:63,122,149-63,122,353 | 1830 | $1.5\times10^{-4}$ | OTX1 | 1.3 |
| | | | | EHBP1 | 3.2 |

[a] ID of topologically associating domain, [b] *p*-value from the HEDD database, [c] disease association score from the DISEASES database.

**Table S3.** Transcription factor (TF) binding sites in physical contact with genetic variants listed in Table S1 according to the 3D genome structure.

| TF | TF DA-score[a] | Genomic position | TAD[b] | TF binding score[c] | Target genes | TG DA-score[a] |
|---|---|---|---|---|---|---|
| | | | *Breast cancer* | | | |
| GATA3 | 5.9 | Chr5:56,109,399-56,111,399 | 6450 | 1090 ($2.2\times10^{-5}$) | MAP3K1 | 5.3 |
| FOXA1 | 5.8 | Chr10:123,353,272-123,355,272 | 12497 | 959 ($4.5\times10^{-4}$) | FGFR2 | 5.6 |
| | | | *Prostate cancer* | | | |
| AR | 8.0 | Chr8:128,426,356-128,428,356 | 8618 | 1005 ($3.5\times10^{-4}$) | POU5F1B | 3.7 |

[a] disease association score from the DISEASES database, [b] ID of topologically associating domain, [c] TF binding score (*p*-value) from the TF2DNA database.
